# Coronary blood vessels from distinct origins converge to equivalent states during mouse and human development

**DOI:** 10.1101/2021.04.25.441380

**Authors:** Ragini Phansalkar, Josephine Krieger, Mingming Zhao, Sai Saroja Kolluru, Robert C. Jones, Stephen R Quake, Irving Weissman, Daniel Bernstein, Virginia D. Winn, Gaetano D’Amato, Kristy Red-Horse

**Affiliations:** Department of Genetics, Stanford University School of Medicine, Stanford, CA, 94305, USA; Department of Biology, Stanford University, Stanford, CA, 94305, USA; Institute for Stem Cell Biology and Regenerative Medicine, Stanford University School of Medicine, Stanford, CA, 94305, USA; Division of Pediatric Cardiology, Department of Pediatrics, Stanford University School of Medicine, Stanford, CA, 94305, UCA; Stanford Cardiovascular Institute, Stanford University School of Medicine, Stanford, CA, 94305, USA; Department of Bioengineering and Department of Applied Physics, Stanford University, Stanford, CA, 94305, USA; Chan Zuckerberg Biohub, Stanford, CA, 94305, USA; Department of Obstetrics and Gynecology, Stanford University School of Medicine, Stanford, CA, 94305, USA

## Abstract

Most cell fate trajectories during development follow a diverging, tree-like branching pattern, but the opposite can occur when distinct progenitors contribute to the same cell type. During this convergent differentiation, it is unknown if cells “remember” their origins transcriptionally or whether this influences cell behavior. Most coronary blood vessels of the heart develop from two different progenitor sources—the endocardium (Endo) and sinus venosus (SV)—but whether transcriptional or functional differences related to origin are retained is unknown. We addressed this by combining lineage tracing with single-cell RNA sequencing (scRNAseq) in embryonic and adult mouse hearts. Shortly after coronary development begins, capillary ECs transcriptionally segregated into two states that retained progenitor-specific gene expression. Later in development, when the coronary vasculature is well-established but still remodeling, capillary Ecs again segregated into two populations, but transcriptional differences were primarily related to tissue localization rather than lineage. Specifically, ECs in the heart septum expressed genes indicative of increased local hypoxia and decreased blood flow. Adult capillary ECs were more homogeneous with respect to both lineage and location. In agreement, SV- and Endo-derived ECs in adult hearts displayed similar responses to injury. Finally, scRNAseq of developing human coronary vessels indicated that the human heart followed similar principles. Thus, over the course of development, transcriptional heterogeneity in coronary ECs is first influenced by lineage, then by location, until heterogeneity declines in the homeostatic adult heart. These results highlight the plasticity of ECs during development, and the validity of the mouse as a model for human coronary development.

## Introduction

During embryonic development, progenitor tissue sources produce new cell types through shifts in epigenetic and transcriptional states. Much research addresses how new cell types form, yet there is less focus on how the transcriptional or chromatin states of progenitor cells relate to gene expression in their descendants, i.e. what do mature cells “remember” about their history? This question is particularly intriguing in cases where multiple progenitor sources contribute to the same cell type, since different origins could result in different behaviors or responses to injury and disease. Such lineage merging is referred to as “convergent differentiation” and occurs in hematopoietic populations (Sathe et al., 2013; Weinreb et al., 2020), oligodendrocytes (Marques et al., 2018), olfactory projection neurons (Li et al., 2017), coronary blood vessels of the heart (Sharma et al., 2017), and others (Wei et al., 2015; Gerber et al., 2018; Konstantinides et al., 2018). Recent single-cell RNA sequencing (scRNAseq) analyses have suggested that the resulting cell types can converge transcriptionally (Li et al., 2017), but in some cases maintain molecular signatures of their progenitors (Dick et al., 2019; Weinreb et al., 2020). However, there is no information on how convergent differentiation influences coronary blood vessels of the heart or how this might affect cardiac injury responses.

In this study, we investigated gene expression patterns in two lineage trajectories that form the coronary vasculature in mice and compared these with data from human fetal hearts. The major progenitor sources for coronary endothelial cells (ECs) in mice are the sinus venosus (SV), the venous inflow tract of the developing heart, and the endocardium (Endo), the inner lining of the heart ventricles (Red-Horse et al., 2010; Wu et al., 2012; Chen et al., 2014; Tian et al., 2014; Zhang et al., 2016; Sharma et al., 2017; Su et al., 2018). ECs begin to migrate from both these sources at embryonic day 11.5. They form an immature capillary plexus (Zeini et al., 2009; Red-Horse et al., 2010) by populating the heart with vessels from the outside-in (SV) or the inside-out (Endo). These two sources eventually localize to largely complimentary regions in adults: the SV contributes vessels to the outer myocardial wall and the Endo contributes vessels to the inner myocardial wall and the septum (Red-Horse et al., 2010; Wu et al., 2012; Chen et al., 2014; He et al., 2014; Tian et al., 2014; Zhang et al., 2016; Sharma et al., 2017). Initial angiogenesis from the SV or Endo is guided by different signaling factors (Wu et al., 2012; Arita et al., 2014; Chen et al., 2014; Su et al., 2018; Payne et al., 2019). However, in circumstances where SV angiogenesis is stunted, Endo-derived vasculature can expand into the outer wall to compensate for the vessel loss (Sharma et al., 2017).

In contrast to the well-characterized spatial differences between Endo and SV angiogenesis, there have been no comparisons of transcriptional states between Endo- and SV-derived coronary vessels in development or in adulthood. In addition, there is little information on whether coronary ECs in humans are also derived from these sources, or whether the transcriptional and functional states that human coronary ECs pass through during development match those in the mouse. Understanding any lineage-specific or species-specific traits would have important implications for approaches that reactivate developmental pathways to increase angiogenesis in injured or diseased human hearts (Smart, 2017; Payne et al., 2019).

Here, we used scRNAseq of lineage-traced ECs from mouse hearts at various stages of development to show that while SV- and Endo-derived capillary cells initially retained some source-specific gene expression patterns, these differences were only present at an early stage of development. Later, these lineages mixed into two capillary subtypes, which were correlated with different locations in the heart. By adult stages, SV- and Endo-derived capillary cells had converged into similar gene expression patterns, and differing lineage did not result in differential proliferation in response to ischemia/reperfusion (IR)-induced injury. Finally, scRNAseq on human fetal hearts indicated that human development closely matched that in mice, and provided additional insights into human coronary artery development. Based on our results, we propose a model in which the transcriptional state of non-proliferative coronary ECs is initially influenced by their lineage, then by regional differences in environmental factors, until both of these signatures fade in adulthood. These findings highlight the importance of environmental factors in influencing EC behavior, and validate mice as a representative model for human coronary development.

## Results

### ScRNAseq in lineage-labeled coronary ECs

To compare Endo- and SV-derived ECs during development and in adult hearts, we combined scRNAseq with lineage-specific fluorescent labeling (Fig. 1a and b). A tamoxifen-inducible *BmxCreER* (Ehling et al., 2013) mouse was crossed with the *Rosa^tdTomato^* Cre reporter, which specifically labels a high percentage of the Endo (94.44% of Endo cells labeled at e12.5), but does not mark the SV (3.61% of SV cells labeled at e12.5)(D’Amato et al., *in preparation*). Labeling was induced before e11.5, when coronary development begins, so Endo-derived ECs expressed *tdTomato* while SV-derived ECs did not (Methods). Cells from e12, e17.5, and adult hearts were sorted using fluorescence-activated cell sorting (FACS) and processed using the 10X Genomics platform (Fig. 1a-b and S1a-e). These strategies captured the expected EC subtypes at each stage (including coronary, valve, Endo, and SV)(Fig. S2a-d) and contained a large number of cells that passed standard quality controls (Methods). ScRNAseq analyses are most accurate when the specific cell populations of interest are extracted and re-analyzed without the influence of other cell types in the dataset, including cycling cells (Luecken and Theis, 2019). Thus, we isolated non-cycling coronary ECs (for e12—*Pecam1+, Cldn5+, Npr3-, Top2a-, Mki67, Tbx20-, Cldn11-, Bmp4-, Vwf-*; for e17.5—*Pecam1+, Npr3-, Tbx20-, Pdgfra-, Top2a-, Bmp4-, Mki67-*)(Fig. S2e and f) and performed direct comparisons of cell states between Endo- and SV-enriched coronary ECs. The remaining cells in the dataset will be reported by D’Amato et al., which addresses experimental questions outside the scope of this study.

**Figure 1:**
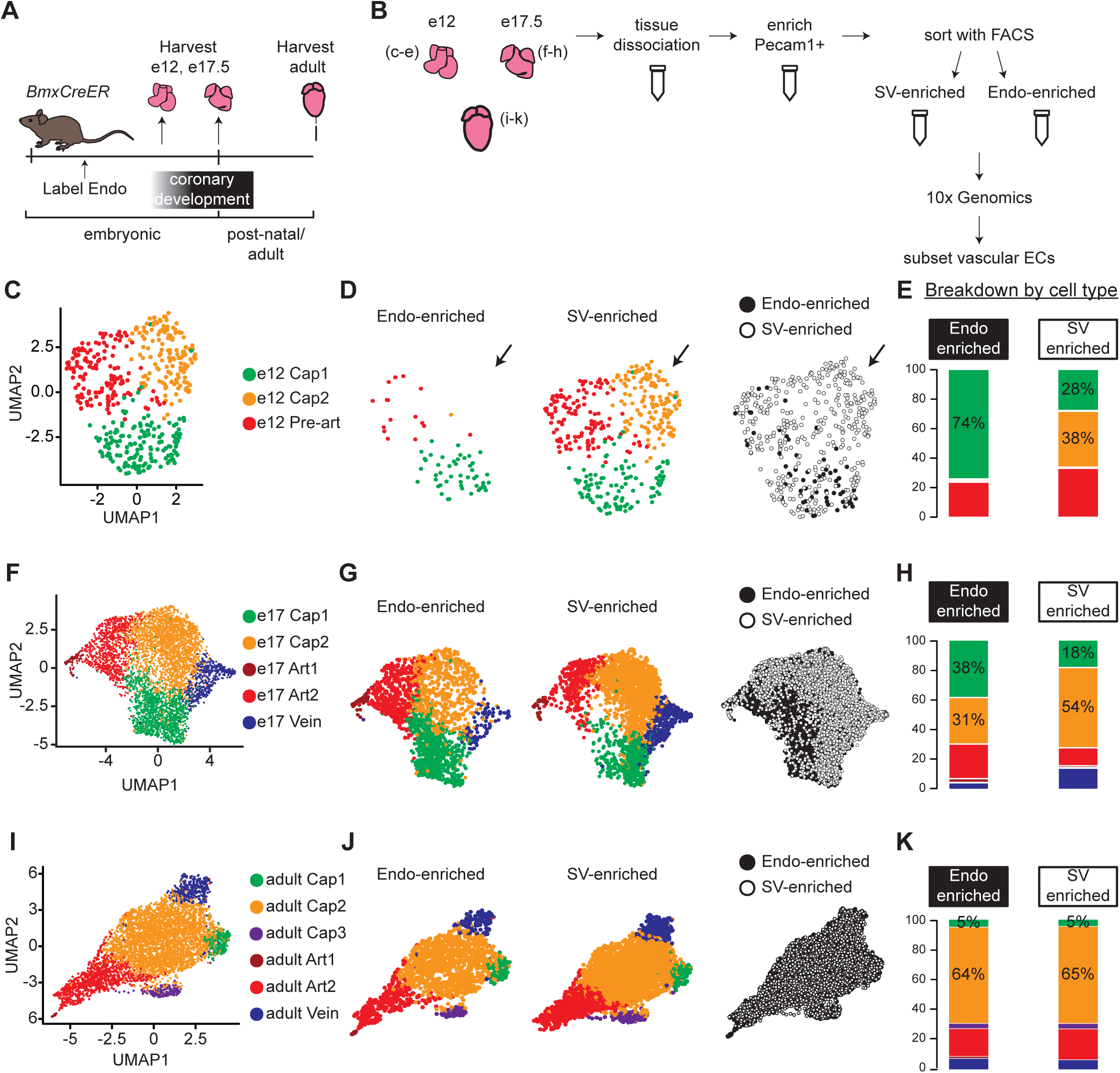
ScRNAseq of lineage-traced coronary ECs at three stages reveals capillary heterogeneity during embryonic development. (A and B) Overview of lineage tracing and scRNAseq approach in embryonic and adult mice. (C-K) Unbiased clustering of embryonic coronary ECs at the indicated time points and the contribution of Endo-enriched (*BmxCreER* lineage-labeled) and SV-enriched (*BmxCreER* lineage negative) cells to each cluster. UMAPs are shown for combined data (C, F, and I) and separated by lineage (D, G, and J) and percentages enumerated (E, H, and K).

We first used unbiased clustering to identify coronary EC subtypes within the e12, e17.5, and adult datasets. Clustering resolution was determined individually for each dataset as the highest resolution at which every cluster expressed at least one unique marker gene. E12 coronary ECs separated into three clusters—capillary plexus 1 (Cap1), capillary plexus 2 (Cap2), and pre-artery (Fig. 1c). Markers used to identify these populations matched previous reports and are shown in Fig. S3a. The absence of venous ECs and the higher numbers of SV-enriched cells present at this stage are also consistent with previous studies (Red-Horse et al., 2010; Su et al., 2018). Separating plots by sample revealed that Cap2 was exclusively from the SV-enriched sample (Fig. 1d and e). Consistent with this was its increased expression of *Apj* (*Aplnr*)(Fig. S3a), which we previously demonstrated to be enriched in SV-derived vessels (Sharma et al., 2017). All but one of the Endo-enriched capillary cells were in Cap1, but Cap1 also contained cells from the SV-enriched sample (Fig. 1d and e). Although recombination rates in the Endo were very high (Methods), we cannot exclude the possibility that a small number of Endo-derived ECs are *tdTomato*-negative due to some un-recombined Endo ECs. These data show that shortly after coronary development is initiated, lineage is correlated with transcriptionally distinct capillary populations within the immature capillary plexus.

To test whether this phenomenon persists into late development, we similarly analyzed coronary ECs at e17.5. A larger number of coronary ECs were captured due to the increase in cardiac vasculature by this stage. EC clusters in this sample included two artery (Art1 and Art2), one vein, and two capillary (Cap1 and Cap2) clusters (Fig. 1f and S3b). If Cap1 and Cap2 continued to reflect different lineages, we would expect at least one cluster to contain only Endo- or SV-enriched ECs. There was skewed contribution with a higher percentage of Endo-enriched cells in Cap1 and a higher percentage of SV-enriched cells in Cap2 (Fig. 1g and h), but no cluster was lineage exclusive, suggesting that additional factors were driving transcriptional heterogeneity. Veins were much more represented in the SV-enriched sample (Fig. 1h), which is expected since most veins reside on the surface of the heart and SV angiogenesis progresses outside-in while Endo angiogenesis is in the opposite direction. We also considered the e17.5 dataset with cycling cells included, but with cell cycle effects regressed out, in order to evaluate whether there are differences in proliferation between Endo- and SV-enriched cells, and to rule this out as a cause for the difference in cluster distribution between the two lineages (Fig. S4a and b). This analysis showed that cycling capillary cells also segregate into the Cap1 and Cap2 clusters (Fig. S4c), and that their distribution into these clusters is biased by lineage, similar to non-cycling cells (Fig. S4d). Additionally, there is no difference in proliferation between Cap1 and Cap2 (Fig. S4e).

We next performed the same analyses on adult coronary ECs. Similarly to e17.5, clustering revealed one vein and two artery clusters. It additionally revealed three capillary clusters, Cap1, Cap2 and Cap3 (Fig. 1i), which were distinguishable by expression of unique gene markers (Fig. S3c and S5a). While Cap1 contained the majority of capillary cells, Cap2 was distinguished by expression of pro-angiogenic genes including *Apln* and *Adm*, and Cap3 was distinguished by expression of interferon-induced genes such as *Ifit3* (Fig. S5a). Both of these clusters have been reported previously in adult hearts (Kalucka et al., 2020). However, unlike the clusters at earlier stages, there was a similar distribution of cells into each of these clusters in both the Endo- and SV-enriched samples (Fig. 1j-k), and there were no appreciable gene expression differences between the samples. This observation is consistent with another lineage-specific adult scRNAseq dataset we produced with Smart-seq2 (unpublished results). Thus, we concluded that there is no lineage-based heterogeneity in adult coronary ECs.

### Coronary heterogeneity is first related to lineage and then to location

We next investigated the genes driving coronary ECs into two capillary cell states at the different stages of development. One hypothesis was that cells retained gene expression patterns from their progenitors. To test this, a list of genes defining the progenitor states (the Endo and SV) was compiled by directly comparing gene expression in the Endo and the SV at e12 (Fig. S2a) and using differentially expressed genes (DEGs) passing significance thresholds described in the *Methods* (Table 1). The expression of these genes was then assessed in capillary clusters. At e12, Cap1 cells expressed higher levels of Endo-specific genes, while Cap2 cells expressed higher levels of SV-specific genes (Fig. 2a and S6a-c). Indeed, 40% of Cap1 genes and 3% of Cap2 genes overlapped with the Endo, while 47% of Cap2 genes and 1% of Cap1 genes overlapped with the SV (Fig. 2b). We concluded that the transcriptional identities of Cap1 and Cap2 cells derive at least in part by gene expression patterns retained from the SV or Endo.

**Figure 2:**
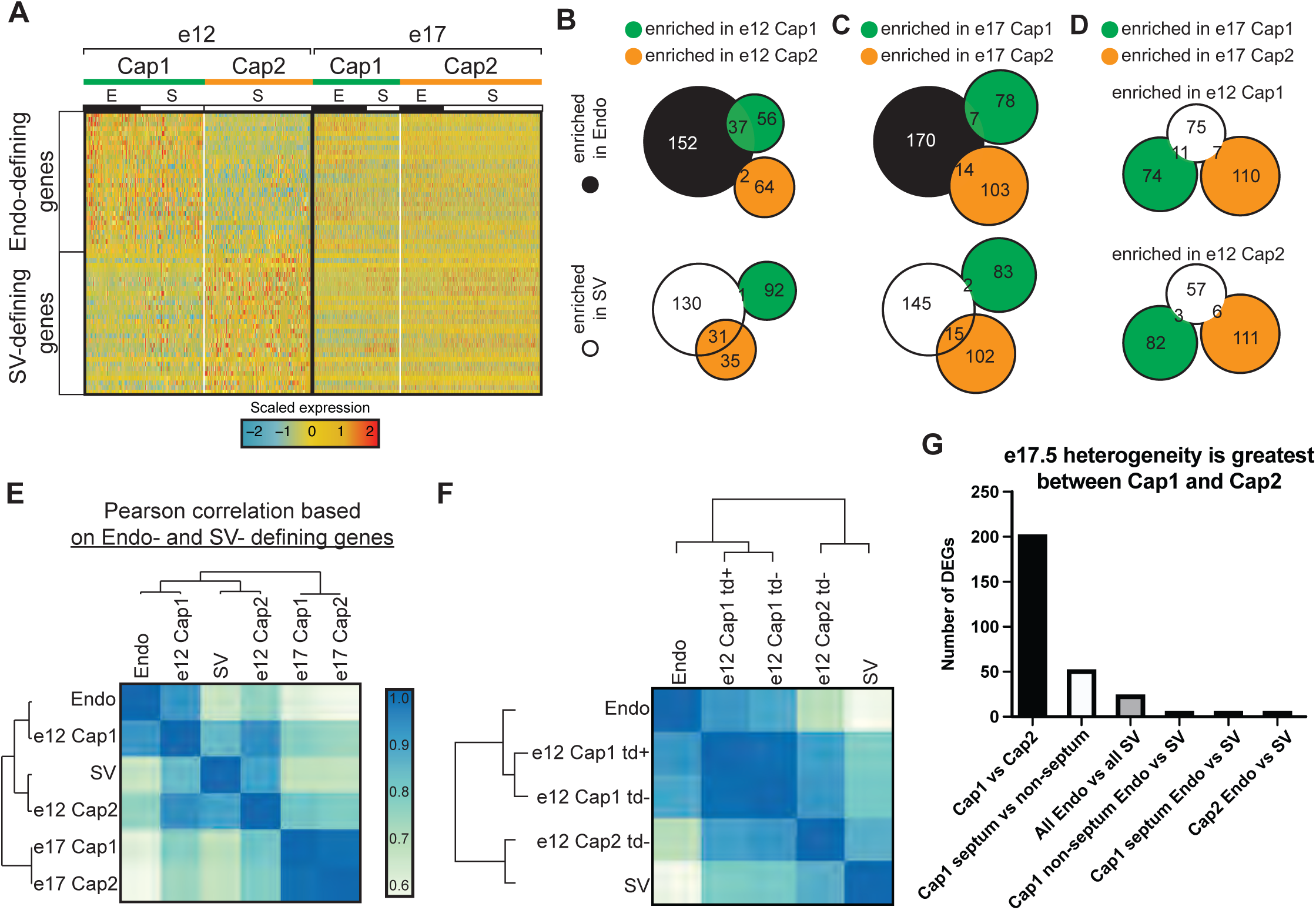
Expression of Endo and SV genes in coronary ECs. (A) Heatmap showing expression of the top 30 (by p-value) Endo-defining genes (enriched in the Endo compared to the SV) and the top 30 (by p-value) SV-defining genes (enriched in the SV compared to the Endo) in e12 and e17.5 capillary clusters (E = coronary cells from the Endo-enriched sample, S = coronary cells from the SV-enriched sample). (B and C) Venn-diagrams showing overlap of Endo- and SV- defining genes with Cap1 enriched genes (enriched in Cap1 compared to Cap2) and e12 Cap2 enriched genes (enriched in Cap2 compared to Cap1) at e12 (B) and e17.5 (C). (D) Venn-diagram showing overlap of e12 Cap1 and Cap2 enriched genes with e17.5 Cap1 and Cap2 genes. (E and F) Heatmaps of Pearson correlations based on expression of Endo- and SV-defining genes in the Endo, the SV, and capillary clusters from e12 and e17.5 in total (E) and separated by *BmxCreER* lineage as indicated by *tdTomato* (*td*) expression (F). (G) Bar plot showing number of DEGs between different subgroups of capillary cells.

**Table 1.** Differentially expressed genes between Endo and SV (from heatmap in Fig 2a).

| **Endo-specific genes** | **SV-specific genes** |
| --- | --- |
| Cdkn1c | Cldn11 |
| Tmem108 | Aplnr |
| Nrk | Agr2 |
| Fabp5 | Bmp4 |
| Cdh11 | Hoxa5 |
| Cd81 | Gm13889 |
| Irx5 | Zfp503 |
| H19 | Hotairm1 |
| Hand2 | Rassf9 |
| Adgrg6 | Mmrn1 |
| Maged2 | Pcdh17 |
| Ccnd2 | Tox |
| Plvap | Slco3a1 |
| Igf2 | Tspan13 |
| Dok | Nr2f2 |
| Col13a1 | Fst |
| Tm4sf1 | Cd36 |
| Igf2r | Cldn5 |
| Gm1673 | Aqp1 |
| Blvrb | Hoxb4 |
| Gsta4 | Khdrbs3 |
| Rap2a | Kitl |
| Gatm | Wnt16 |
| Olfml3 | Limch1 |
| Sdpr | Ahr |
| Ece1 | Edn1 |
| Plagl1 | Lamp5 |
| Prr15 | Cav1 |
| Igfbp4 | Ddah1 |
| Ccnd3 | Tbx5 |

This pattern was not observed at e17.5. There were no clear pattern between e17.5 Cap1 and Cap2 in the expression of SV- and Endo-specific genes (Fig. 2a and S6d), and there was little to no overlap between Cap1 and Cap2 differential genes and SV or Endo genes (Fig. 2c). Although it may appear on the heatmap in Fig. 2a that there is a lineage-based distinction in Cap1 for a small subset of the genes at e17.5, this is not the case. As we will demonstrate later in Fig. S9, the data indicated that these results were primarily due to Cap1 containing septum ECs, which are mostly derived from the Endo, and the septum imparting a location-specific effect on transcription. Furthermore, the minimal overlap between the e12 and e17.5 Cap1- and Cap2-defining genes is consistent with e17.5 coronary ECs not retaining the progenitor-type genes enriched in e12 clusters (Fig. 2d).

Calculating Pearson correlations using Endo and SV genes revealed that e12 coronary ECs were similar to their progenitor sources while e17.5 coronary ECs were much less so (Fig. 2e).

We also calculated Pearson correlations as a function of lineage at e12. As expected, *tdTomato* positive cells were highly similar to the Endo while *tdTomato* negative cells in Cap2 were more similar to the SV than to the Endo (Fig. 2f). Interestingly, *tdTomato* negative cells in Cap1 were more similar to the Endo than to the SV (Fig. 2f). This could result from a number of reasons that we cannot currently distinguish including: 1) SV-derived cells migrating close to the Endo take on Endo-type gene expression or 2) There is a rare Endo population that does not express *Bmx*. Point 2 is supported over point 1 due to the observation that Cap2-like cells are present in a previous scRNAseq dataset of SV-derived ECs (Su et al., 2018), while Cap1 cells are not (Fig. S7a-e). In total, these data indicate that e17.5 capillary heterogeneity is not a remnant of the heterogeneity present at e12, and lineage-related differences are not apparent in adults.

Further analyses provided additional evidence that the differences between e17.5 Cap1 and Cap2 are not primarily due to lineage. With the prediction that significant lineage-related heterogeneity would be accompanied by substantial differences in gene expression, we compared the DEGs between e17.5 Cap1 and Cap2 to the DEGs between Endo- and SV-enriched capillary cells. There were 202 DEGs between e17.5 Cap1 and Cap2, but only 24 DEGs between all Endo enriched and SV-enriched capillaries (Fig. 2h). Inspecting DEG identities provided further support that lineage is not retained. 18 of the DEGs between all Endo- and SV-enriched cells were also DEGs between Cap1 and Cap2 (Table 2). If the differential patterns of these genes were due to a lineage effect, we would expect a greater log-fold change in the all Endo- versus SV-enriched comparison than in the Cap1 versus Cap2 comparison. However, 16 of the 18 genes have a greater log fold change in the Cap1 versus Cap2 comparison (Table 2). These data indicate that differential expression between the Endo- and SV-enriched capillaries mostly stems from the differential contribution of the Endo and SV lineages to Cap1 and Cap2.

**Table 2.** Differentially expressed genes between all Endo-enriched and all SV-enriched capillaries. Bolded genes are also differentially expressed between Cap1 and Cap2.

|  | Raw p-value | Average log-fold change | % Endo cells expressing | % SV cells expressing | Adjusted p-value | Higher fold change in Cap1 vs Cap2 comparison? |
| --- | --- | --- | --- | --- | --- | --- |
| tdTomato | 1.91E-238 | 0.39378511 | 0.452 | 0 | 5.35E-234 |  |
| Gt(ROSA)26Sor | 2.43E-153 | -0.634801942 | 0.568 | 0.837 | 6.81E-149 |  |
| **Anxa1** | 4.24E-109 | 0.553933954 | 0.804 | 0.53 | 1.19E-104 |  |
| Igf2 | 6.74E-93 | -0.357191579 | 0.99 | 0.999 | 1.89E-88 |  |
| **Timp4** | 1.28E-88 | -0.632710228 | 0.447 | 0.766 | 3.58E-84 | yes |
| **Fmo1** | 8.34E-70 | -0.351754806 | 0.231 | 0.511 | 2.34E-65 | yes |
| **Txnip** | 2.09E-67 | -0.37131915 | 0.851 | 0.933 | 5.85E-63 | yes |
| **Aplnr** | 5.42E-67 | -0.40125981 | 0.677 | 0.863 | 1.52E-62 | yes |
| **Car4** | 1.68E-65 | -0.595662824 | 0.347 | 0.62 | 4.70E-61 | yes |
| **Aqp7** | 3.33E-62 | -0.368823615 | 0.345 | 0.63 | 9.31E-58 | yes |
| Apoe | 1.19E-60 | 0.302862096 | 0.307 | 0.09 | 3.34E-56 |  |
| **Cd36** | 8.58E-59 | -0.308565577 | 0.97 | 0.997 | 2.40E-54 | yes |
| **Igfbp5** | 1.84E-57 | 0.4602195 | 0.258 | 0.064 | 5.15E-53 |  |
| **Fabp5** | 6.53E-54 | 0.30861622 | 0.999 | 1 | 1.83E-49 | yes |
| **1810011O10Rik** | 1.08E-52 | -0.340249285 | 0.907 | 0.963 | 3.04E-48 | yes |
| **Aqp1** | 9.79E-47 | -0.471913791 | 0.54 | 0.729 | 2.74E-42 | yes |
| **Gap43** | 7.23E-38 | 0.34418651 | 0.523 | 0.331 | 2.02E-33 | yes |
| **Sat1** | 1.99E-37 | 0.394143816 | 0.753 | 0.63 | 5.59E-33 | yes |
| **Cd63** | 3.81E-36 | 0.33008301 | 0.758 | 0.611 | 1.07E-31 | yes |
| **Igfbp3** | 3.45E-34 | 0.640208849 | 0.634 | 0.468 | 9.66E-30 | yes |
| Tm4sf1 | 1.38E-32 | 0.302207793 | 0.982 | 0.94 | 3.87E-28 |  |
| Maged2 | 4.03E-32 | 0.323885527 | 0.845 | 0.757 | 1.13E-27 |  |
| **Ly6c1** | 1.32E-31 | -0.301696479 | 0.845 | 0.93 | 3.70E-27 | yes |
| **Rbp1** | 1.26E-15 | 0.432393695 | 0.757 | 0.69 | 3.52E-11 | yes |

Since there was not strong evidence that lineage was a significant factor, we next considered whether differential localization in the heart might underlie e17.5 heterogeneity. During development, different regions of the heart show varying levels of hypoxia and signaling factors, e.g. at some stages, the septum is more hypoxic and expresses higher levels of *Vegfa* (Miquerol et al., 2000; Sharma et al., 2017). Localization-driven heterogeneity would also explain the bias in cluster distribution between the Endo and SV lineages (Fig. 1h) because they contribute ECs to complementary regions of the heart. This is most dramatic in the septum where almost all ECs are derived from the Endo (Wu et al., 2012; Chen et al., 2014; Zhang et al., 2016). The top DEGs between e17.5 Cap1 and Cap2 included hypoxia-induced genes (*Mif, Adm, Igfbp3, Kcne3*)(Tazuke et al., 1998; Lee et al., 1999; Keleg et al., 2007; Simons et al., 2011; Heng et al., 2019) and tip cell markers (*Apln, Plaur, Lamb1, Dll4*)(Hellstrom et al., 2007; del Toro et al., 2010) in Cap1, and flow-induced genes (*Klf2, Klf4, Thbd, Lims*) (Dekker et al., 2002; Hamik et al., 2007; Kumar et al., 2014; Wang and Zhang, 2020) in Cap2 (Fig. 3a and b, Fig. S8a), suggesting pathways that could be consistent with localization-driven heterogeneity.

**Figure 3:**
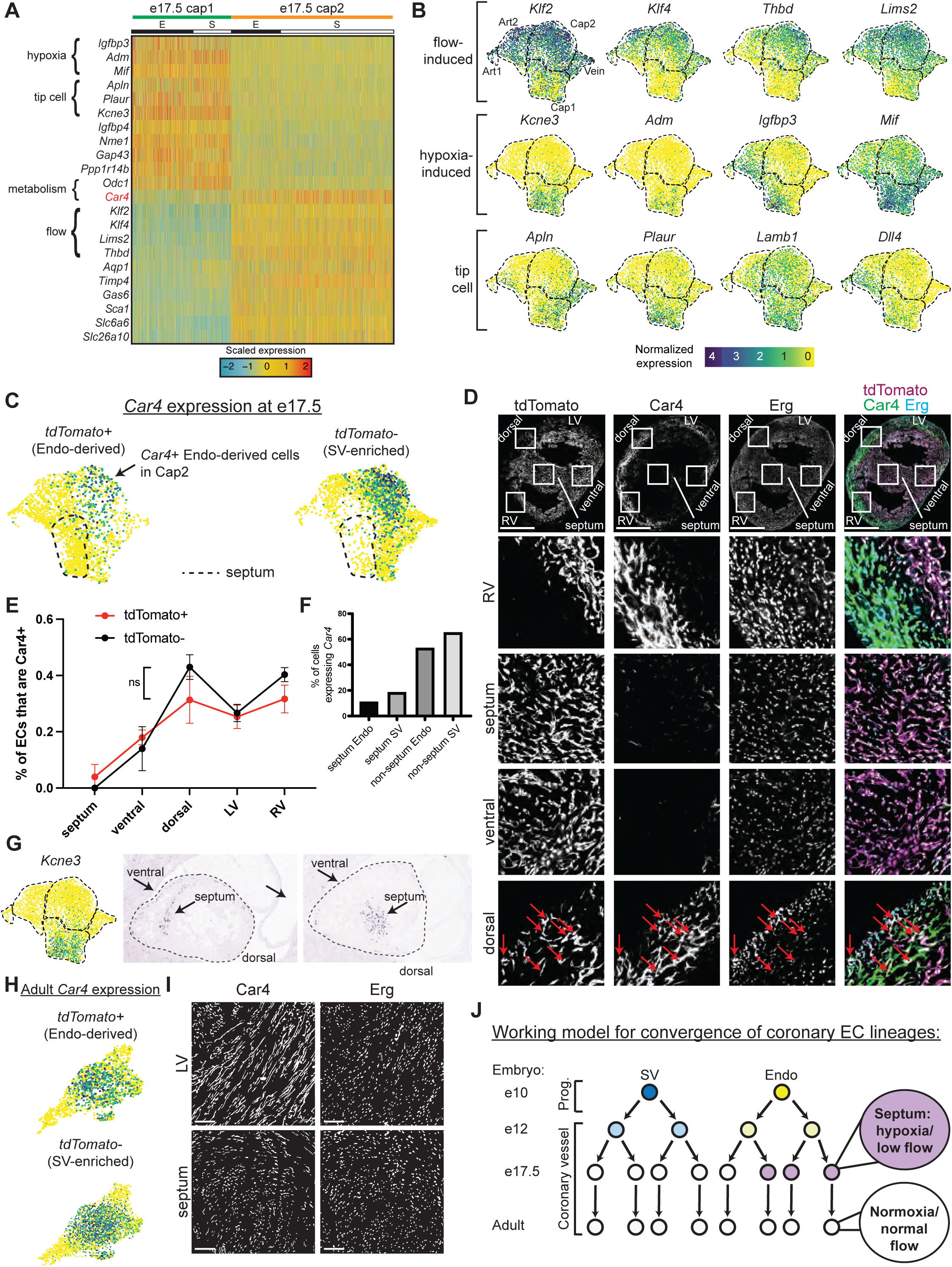
Gene expression and localization of e17.5 capillary clusters. (A) Heatmap showing expression of selected genes enriched in either Cap1 or Cap2 at e17.5 (E = coronary cells from Endo-enriched sample, S = coronary cells from SV-enriched sample). (B) UMAPs showing expression of selected flow-induced, hypoxia-induced, and tip-cell genes. Dashed lines outline indicated clusters. (C) *Car4* UMAPs separated by lineage. Dashed line shows area of UMAP enriched in Endo-enriched, *Car4*-negative cells predicted to be located in the septum. (D) Immunofluorescence of Car4 and Erg in a heart section from an e17.5 *BmxCreER;Rosa^tdTomato^* embryo (scale bar = 500 μm). Red arrows indicate Car4-positive, tdTomato-positive Endo-derived ECs in the dorsal wall. (E) Plot showing percentage of tdTomato-positive and tdTomato-negative ECs in different locations which are also Car4-positive (based on quantification of Car4 staining in Erg-positive cells from three e17.5 *BmxCreER;Rosa^tdTomato^* embryos (error bars = range) (F) Bar plot based on e17.5 scRNAseq showing the percent of capillary cells in different categories (septum Endo-enriched, septum SV-enriched, non-septum Endo-enriched, non-septum SV-enriched) which express *Car4* at any level. (G) Images showing in-situ hybridization for *Kcne3* in an e14.5 embryonic mouse heart, obtained from GenePaint. (H) UMAPs showing expression of *Car4* in adult coronary ECs, separated by lineage. (I) Immunofluorescence of Car4 and Erg in the left ventricle (LV) and septum of an adult WT heart (scale bar = 100 μm). (J) Working hypothesis for convergence of Endo- and SV-derived ECs into equivalent transcriptional states. Scale bar from (B) also applies to (C), (G), and (H).

We next used Car4 to localize Cap2 within the tissue because it was specifically expressed in this cluster and there was an antibody available for immunostaining (Fig. 3a and c). Erg-positive ECs expressing Car4 were located mainly in the right and left ventricle free walls and dorsal side while ECs in the septum and ventral wall were mostly Car4 negative (Fig. 3d-f and S8b). This indicated that Cap1 localizes primarily to the septum and ventral wall and Cap2 to the remaining walls of the ventricle. The *Kcne3* pattern provided further support for this localization because it was specific to Cap1, and *in situ* hybridization revealed specific septal and ventral expression (Fig. 3g). *Car4* analysis revealed a further segregation of Cap1 into septum and non-septum regions on the UMAP. Specifically, one side of the Cap1 cluster was almost completely devoid of *Car4* and comprised almost all Endo-enriched cells (Fig. 3c), which are two features specific to the septum as shown in Fig. 3d-f and S8b. Additionally, these cells have lower expression of *Aplnr* compared to the non-septal cells of Cap1 (Fig. S3b), which is also characteristic of the septum (unpublished observations). Although unbiased clustering did not distinguish this group of cells as a separate cluster, this could be because we used a combination of gene expression, protein staining, and lineage information (Fig. 3b-g), which the clustering algorithms do not take into account. Interestingly, this proposed septum region is where gene expression indicates both decreased blood flow and local hypoxia (Fig. 3b), the former of which would cause the latter. Thus, the coronary vasculature in the septum (and where the rest of Cap1 localizes) may not receive full blood flow at this late stage in development.

With the knowledge that half of Cap1 was almost completely represented by Endo-derived cells from the septum, we revisited the issue of lineage-based heterogeneity by manually searching for genes enriched in the proposed septum and analyzing whether they were correlated with lineage. A few of the Endo-specific genes from the heatmap in Fig. 2a were expressed at a higher level in the septum while a few SV-specific genes were expressed at a lower level (Fig. S9a-d). If this is due to retention of lineage information, we would expect to see the Endo-specific genes expressed in higher percentages of Endo-enriched cells compared to SV-enriched cells both inside and outside the septum. However, there was generally a decrease in the expression of these genes in both Endo- and SV-enriched cells outside of the septum, supporting the location, but not the lineage, hypothesis. Only two genes somewhat followed a pattern indicating lineage retention—*Gucy1b3* and *Hand2*— but these differences did not pass our pre-set significance thresholds (*Methods*) and were expressed in a low percentage of cells (<20% outside the septum) (Fig. S9b).

We performed additional analysis comparing the number of DEGs between Endo- and SV enriched cells within different capillary sub-groups defined by transcriptional states (i.e. clustering) and in different locations, specifically, the proposed septal and non-septal cells of Cap1 and Cap2 (Fig. 2h). If lineage was a major contributor to e17.5 heterogeneity, we would expect to see a substantial number of DEGs between Endo- and SV-enriched cells within a specific location. Instead, once the effect of location was removed by only comparing Endo- and SV-enriched cells either inside or outside of the septum, there were only 6 DEGs between the lineages (Fig. 2h). Further supporting the impact of location in transcription, the second largest number of DEGs was between the septum and non-septum cells of Cap1 (Fig. 2h). Thus, although we cannot exclude that a few genes retain lineage information in a small number of cells, the overwhelming evidence of this analysis supports that location has a greater effect on cell state at e17.5, with little to no contribution by lineage.

To probe whether the septal portion of Cap1 (see Fig. 3c, dotted line) segregates based on cell-autonomous, lineage-specific features of Endo-derived ECs or regional environments, we took advantage of the fact that some Endo-lineage labeled cells migrate into SV-biased territories during development (Chen et al., 2014). Cell-autonomous lineage differences would be supported if these cells remained Car4-negative in the ventricular walls. However, the opposite was true. Endo-lineage cells outside the septum were more likely to express Car4 than those in the septum (on average, 23% outside septum vs. 4% inside septum)(Fig. 3f, Fig. S8b). Endo-derived ECs in SV-biased territories such as the dorsal side of the heart start to express Car4 (Fig. 3d, arrows), and they can exist in the Cap2 state (Fig. 3c, arrow). In addition, the rates of Car4 positivity in both Endo- and SV-enriched ECs correlate similarly with location in the heart (Fig 3e). Thus, the combination of lineage labeling, scRNAseq, and histology allowed us to ascertain that Endo- lineage ECs are biased towards a separate cell state based on their preferential location in hypoxic regions. However, due to the strong association between EC lineage and location, we cannot rule out that some minor degree of lineage-based heterogeneity exists at e17.5.

Since regional hypoxia would be incompatible with adult heart function, we predicted that the absence of strong heterogeneity in adult capillary ECs could be explained by a resolution of regional environmental differences after development. To test this, we examined *Car4* in the adult dataset, and found that it is broadly expressed in all adult capillary populations (Fig. 3h). Additionally, immunostaining confirmed that septum ECs had become positive for Car4 in adults (Fig. 3i, Table 3). Altogether, our data support the following model—that over the course of development, transcriptional heterogeneity in coronary ECs is first influenced by lineage, then by location, until both lineage- and location-based heterogeneity disappear in the static adult heart (Fig. 3j).

**Table 3.** Ratio of Erg+Car4+/Erg+ cells in adult mouse hearts.

|  | LV | Septum |
| --- | --- | --- |
| heart1 | 0.874 | 0.853 |
| heart2 | 0.864 | 0.819 |

### Lineage does not change response to ischemia-reperfusion injury

Although Endo- and SV-derived capillary ECs were transcriptionally very similar in normal adult hearts, they could behave differently when challenged with hypoxia or other injury. To test this, we performed an ischemia-reperfusion (IR) injury by temporarily occluding the left anterior descending (LAD) coronary artery in adult *BmxCreER;Rosa^tdTomato^* mouse hearts in which the Endo was labelled prior to coronary development (Fig. 4a and S1b). Proliferation in Endo- and SV-enriched ECs was quantified using EdU incorporation assays on day 5 post-injury. This time point was chosen based on prior data showing regrowth of the majority of coronary vessels 5 days after IR injury (Merz et al., 2019). Observing overall EdU incorporation revealed the site of injury, which was prominent in the mid-myocardial region between the inner and outer wall (Fig. 4b). Proliferation was assessed in areas just below the ligation and apex of the heart, as this is the expected distribution of ischemia following LAD ligation (Merz et al., 2019). Regions of interest were chosen in myocardial areas containing a mix of Endo- and SV-derived ECs to ensure adequate representation from both lineages (Fig. 4c and d). This analysis found no difference between the lineages (Fig. 4e). We next performed EdU quantification in the inner and outer walls of the myocardium where most ECs derive from either the Endo or SV, respectively (Wu et al., 2012; Tian et al., 2014; Zhang et al., 2016; Sharma et al., 2017)(Fig. 4b and S1b). In contrast to the lineage comparison, the outer wall showed a consistent, though non-significant, increase in EC proliferation over the inner wall, both within and adjacent to the injury site and regardless of lineage, indicating that this injury and proliferation assay can reveal differences (Fig. 4f). These data support the notion that the local environment, rather than lineage, regulates capillary responses to injury in adult hearts, at least with respect to EC proliferation in the days after injury.

**Figure 4:**
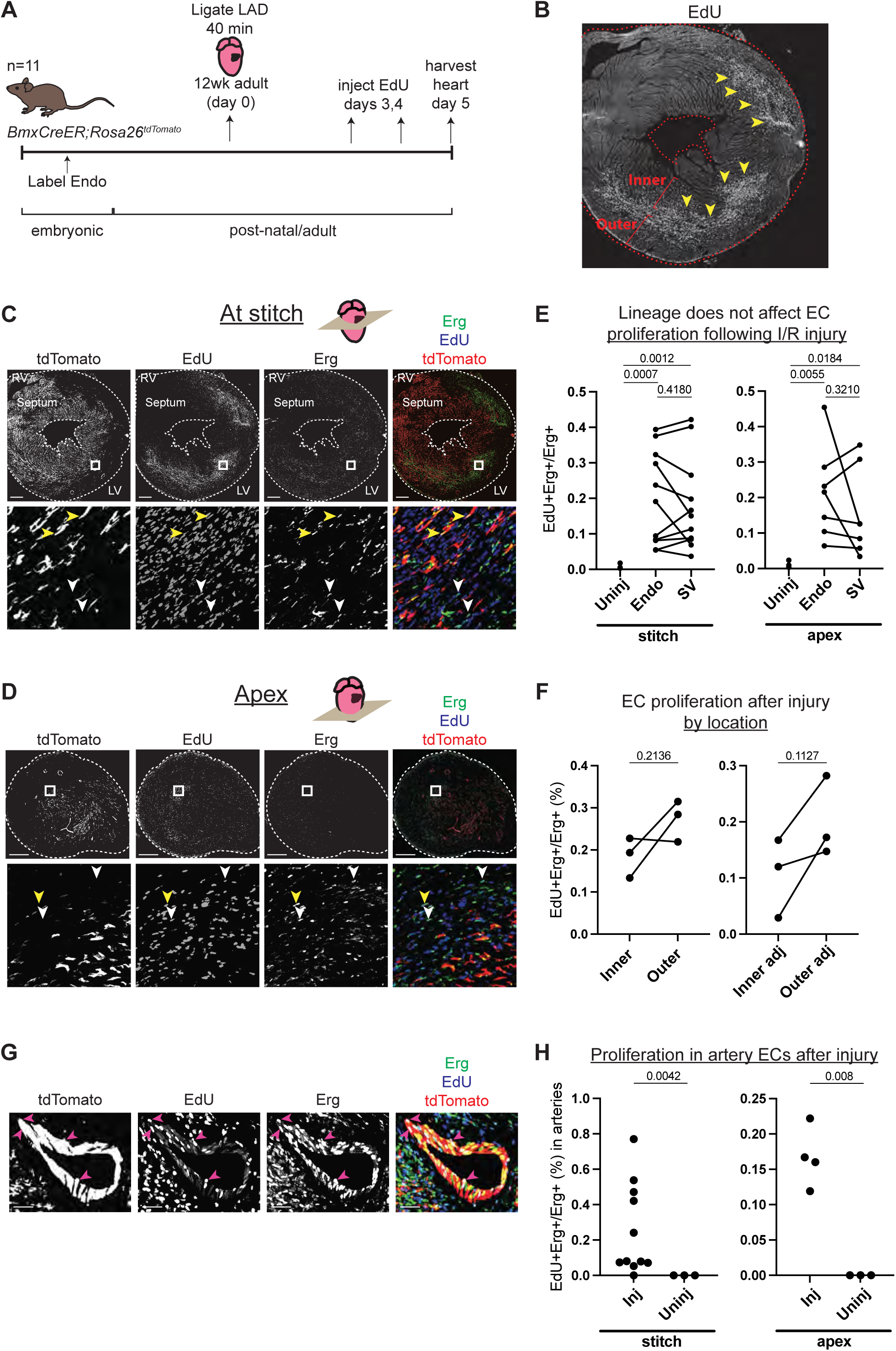
Comparison of injury responses of Endo- and SV-derived coronary ECs. (A) Overview of lineage tracing and ischemia-reperfusion (I/R) injury approach in adult mice. (B) Example of how EdU localization highlights mid-myocardial injury region. Yellow arrowheads indicate the injury region with dense EdU staining. (C and D) Immunofluorescence of EdU and Erg in sections of the heart from (B) just below level of the stitch (C) and in the apex (D). Yellow arrowheads show proliferating Endo-derived ECs that are positive for tdTomato, Erg, and EdU; white arrowheads show tdTomato-negative, Erg-positive ECs from the SV that are EdU positive. (E) Quantification in multiple injured hearts of EdU-positive, Erg-positive ECs from the two lineages. (F) Quantification in multiple injured hearts of EdU-positive, Erg-positive ECs from the inner and outer wall, both in the focal area of the injury, as indicated in (B), and in areas adjacent to the injury. (G) Immunofluorescence of EdU and Erg in an artery of an injured heart. Pink arrowheads show proliferating ECs that are positive for tdTomato, Erg, and EdU. (H) Quantification in multiple injured hearts of EdU-positive, Erg-positive ECs in arteries in the focal area of the injury. In (E), (F), and (H), each dot represents one heart. Scale bar = 50 μm for (G). Scale bars = 500 μm for all other images.

ECs exit the cell cycle during their differentiation into mature coronary arteries (Fang et al., 2017; Su et al., 2018). As a result, the proliferation of artery ECs is rare in the normal adult heart vasculature. Because IR injury induced the proliferation of capillary ECs, we investigated whether IR injury also induced the proliferation of artery ECs (Fig. 4g). Approximately 69% of large arteries (n = 32) identified in the injured regions that we analyzed (as indicated in Fig. 4b) contained at least one EdU-positive EC, and there was a significantly higher rate of EdU-positive ECs in the arteries of injured compared to uninjured hearts (Fig. 4h). This observation could be due either to proliferating capillary cells which transitioned into artery cells but retained EdU, or to artery ECs which began proliferating in response to the injury. Further studies will be necessary to determine whether one or both of these processes are occurring.

### Analogous features in mouse and human coronary ECs

We next sought to investigate whether comparing mouse and human scRNAseq datasets could provide insights into human development. ScRNAseq was performed using Smart-seq2 on PECAM1-positive ECs sorted from human fetal hearts at 11, 14, and 22 weeks of gestation. In addition, a capillary-specific marker, CD36, allowed for enrichment of PECAM1+CD36- arterial ECs (Fig. 5a and S10a and b) (Cui et al., 2019). After initial filtering, 2339 high-quality, high-coverage single EC transcriptomes were obtained of which 713 were arterial. The data included 12 clusters of the expected cell types—artery, capillary, vein, cycling, Endo, valve EC—identified by known markers (Fig. 5b and S10c). As with the mouse data, the analysis was restricted to non-cycling arteries, capillaries, and veins in order to specifically compare cell states and trajectories in coronary vessels. Similar to mouse, the data contained one vein and two capillary clusters, but in contrast to mouse, there was an additional arterial cluster for a total of three (Fig. 5c). There was approximately equal representation of the clusters in the PECAM1-positive fraction at each gestational age, consistent with the data being from later stages (mouse equivalent of e15.5-18.5)(Krishnan et al., 2014) when the developing heart is growing in size rather than producing new coronary cell subtypes (Fig. 5d and S11a). Consequently, we pooled data from the three timepoints for all further analyses. To begin comparing human and mouse, the *Seurat* Label Transfer workflow (Stuart et al., 2019) was utilized to reference map each human cell to its closest mouse cluster and vice versa (Fig. 5e). This showed close concordance between the two species (Fig. 5f and g). With respect to the two capillary clusters that were extensively studied above, the majority of human Cap1 cells mapped to mouse Cap1, while the majority of human Cap2 cells mapped to mouse Cap2 (Fig. 5f). When assigning mouse cells to human clusters, mouse Cap2 almost completely mapped to human Cap2, while mouse Cap1 mapped substantially to human Cap1, Cap2, and Art3 (Fig. 5g). Analyzing specific gene expression revealed several enriched genes shared between corresponding mouse and human capillary clusters (including *KIT*, *ODC1*, *CD300LG*, *RAMP3*), and showed that human Cap1, like its mouse correlate, displayed patterns indicative of experiencing low blood flow conditions and potentially increased hypoxia, i.e. lower *LIMS2, THBD, KLF2, KLF4* and higher *ADM, IGFBP3, KCNE3, LAMB1* (Fig. 5h and S10d). Notably, *CA4* (the human homolog of mouse *Car4*) was not differentially expressed between human Cap1 and Cap2.

**Figure 5:**
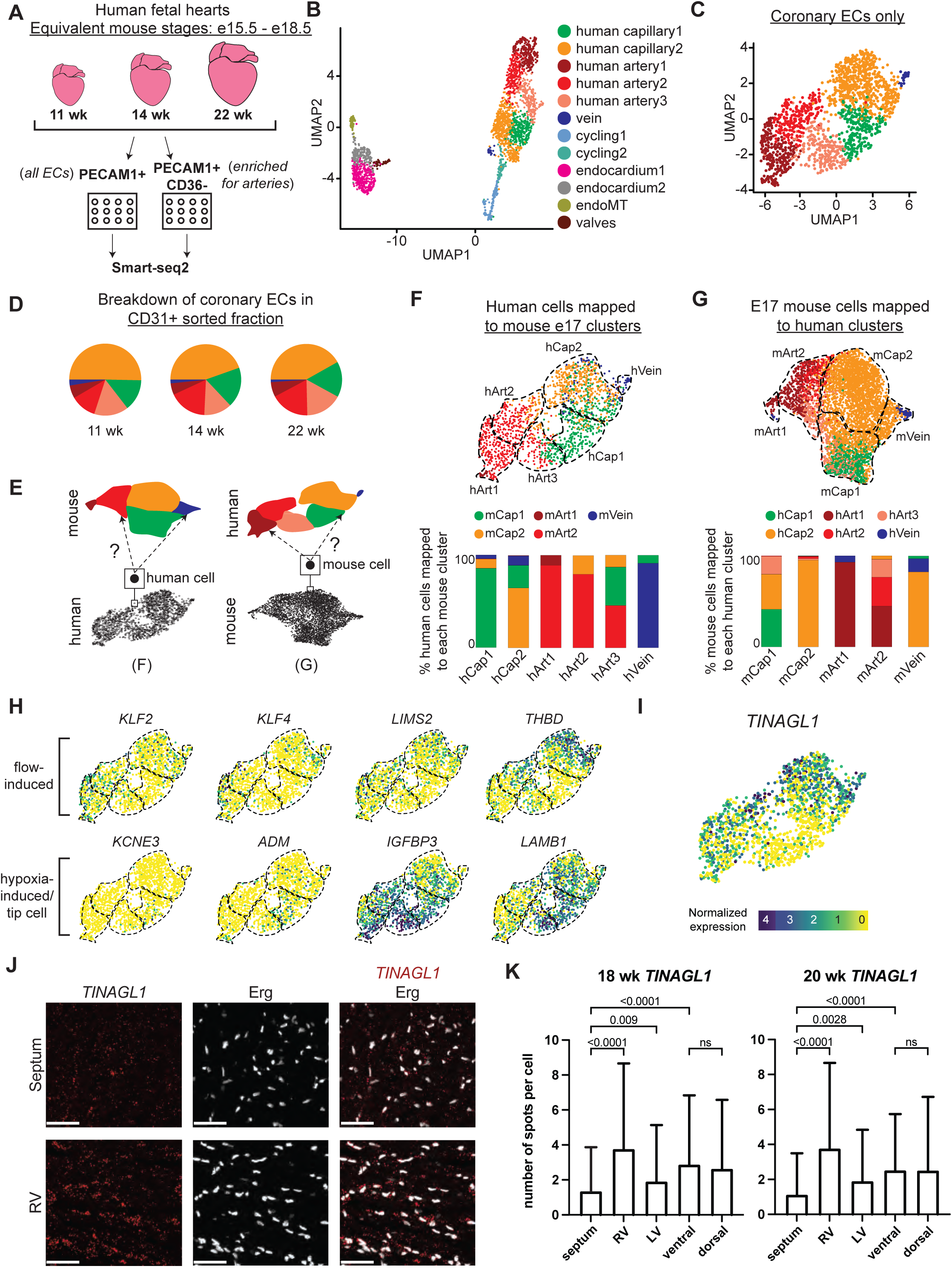
ScRNAseq of coronary ECs from human fetal hearts. (A) Overview of scRNAseq approach for three human fetal hearts. (B and C) UMAPs of all major *PECAM1*+ EC subtypes collected (B) and the non-cycling coronary EC subset (C). (D) Pie charts showing the breakdown by cluster of human coronary ECs that were sorted as *PECAM1*+ without additional enrichment. (E) Schematic of inter-species reference mapping. Individual cells from the human or mouse e17.5 datasets were assigned to the most similar mouse or human cluster, respectively. (F and G) Results from inter-species reference mapping based on shared gene expression, showing the mouse cluster that each human EC mapped to and the percentage breakdown of the mapping from each human cluster (F) and the converse comparison (G). Dashed lines show the borders of the previously defined human and mouse e17.5 coronary clusters. (H) UMAPs showing expression of selected flow-induced, hypoxia-induced and tip-cell genes in human coronary ECs. (I) UMAP showing expression of *TINAGL1* in human coronary ECs. Scale bar from (I) also applies to (H). (J) Section from 18wk human fetal heart showing *in-situ* hybridization for *TINAGL1* with immunofluorescence for Erg. Scale bar = 50 μm. (K) Bar plot showing the mean number of *TINAGL1* RNA spots per cell detected in different regions of 18 wk and 20 wk human fetal hearts. Error bars represent standard deviation.

Knowing that Cap1 and Cap2 segregate spatially in the mouse heart (Fig. 378 3d-f), we examined whether human Cap1 and Cap2 also represent cells in different locations. We performed *in situ* hybridization for *TINAGL1*, a gene which is enriched in human Cap2 (Fig. 5i). This revealed a statistically significant difference in the amount of *TINAGL1* RNA detection between the septum and the heart walls in both an 18 week and a 20 week gestational heart, with the septum having dramatically lower expression (Fig. 5j-k). This difference was especially pronounced between the septum and the right ventricular free wall, similar to the Car4 pattern in mouse (Fig. 3e and S8b). In contrast to mouse Car4, there was no difference in *TINAGL1* detection between the ventral and dorsal walls (Fig. 5k). Thus, *in situ* hybridization with *TINAGL1* supports a bias in the localization of human Cap1 to the septum and human Cap2 to the ventricular wall. Since lineage data and *in situ* immunofluorescence confirmed a subset of mouse Cap1 as containing the Endo-derived cells present in the septum (Fig. 3c), the human expression data in Fig. 5h-k suggested that human Cap1 may also represent an enrichment of septal cells with less blood flow and could also be biased towards the Endo lineage, although the latter cannot be confirmed with gene expression alone. When using the Label Transfer workflow to specifically map the putative mouse septal ECs to the human data, a higher percentage of human Cap1 than Cap2 cells matched to this septal EC group (35% of Cap1 cells versus 14% of Cap2 cells) (Fig. S10e). We concluded that similar developmental environments and endothelial cell states exist between mouse and human capillaries, including those unique to the septum.

### Characterization of the capillary-to-artery transition in human coronary Ecs

Because unbiased clustering produced three artery clusters in human but only two in mouse, we next investigated whether human hearts contained an artery cell state not present in mouse. This could occur because mouse and human arteries have some structural differences, e.g. large conducting arteries in humans are on the surface rather than within the myocardium as in mouse (Wessels and Sedmera, 2003; Kumar et al., 2005; Fernandez et al., 2008; Sorop et al., 2020). If these differences translated into an artery transcriptional state unique to human, we would expect one of the human artery clusters to not be represented in the mouse mapping. Instead, mouse cells mapped to all three human artery clusters (Fig. 6a). There was also evidence indicating that human Art3 cells were in a less mature arterial state compared to human Art1 and 2. This is because: 1) Art3 was unique among the human artery clusters in that a high proportion of Art3 cells matched mouse capillaries (Fig. 6a). 2) Trajectory analysis with RNA velocity (La Manno et al., 2018), Slingshot (Street et al., 2018), and partition-based graph abstraction (PAGA) (Wolf et al., 2019) indicated a transition from human Art3 -> Art2 -> Art1 (Fig. 6b and S11b). and 3) Comparing the fetal dataset with a publicly available adult dataset (Litvinukova et al., 2020) showed that almost no adult human artery ECs mapped to Art3, which would be predicted if Art3 were an immature developmental state (Fig. 6c and S10f). The direction of the arterial trajectory was determined by the arrows from the RNA velocity analysis, which provides this directional information as a consequence of comparing spliced (mature) to unspliced (immature) transcripts (La Manno et al., 2018). Additionally, this trajectory is supported by previous lineage analyses in mouse of a trajectory from capillaries to *Gja5*- arteries to *Gja5*+ arteries (Su et al., 2018). Together, these data suggest that human hearts do not contain a dramatically unique artery transcriptional state when compared to mouse, but they do have an immature state (Art3) that matures into Art1 and Art2 in adults.

**Figure 6:**
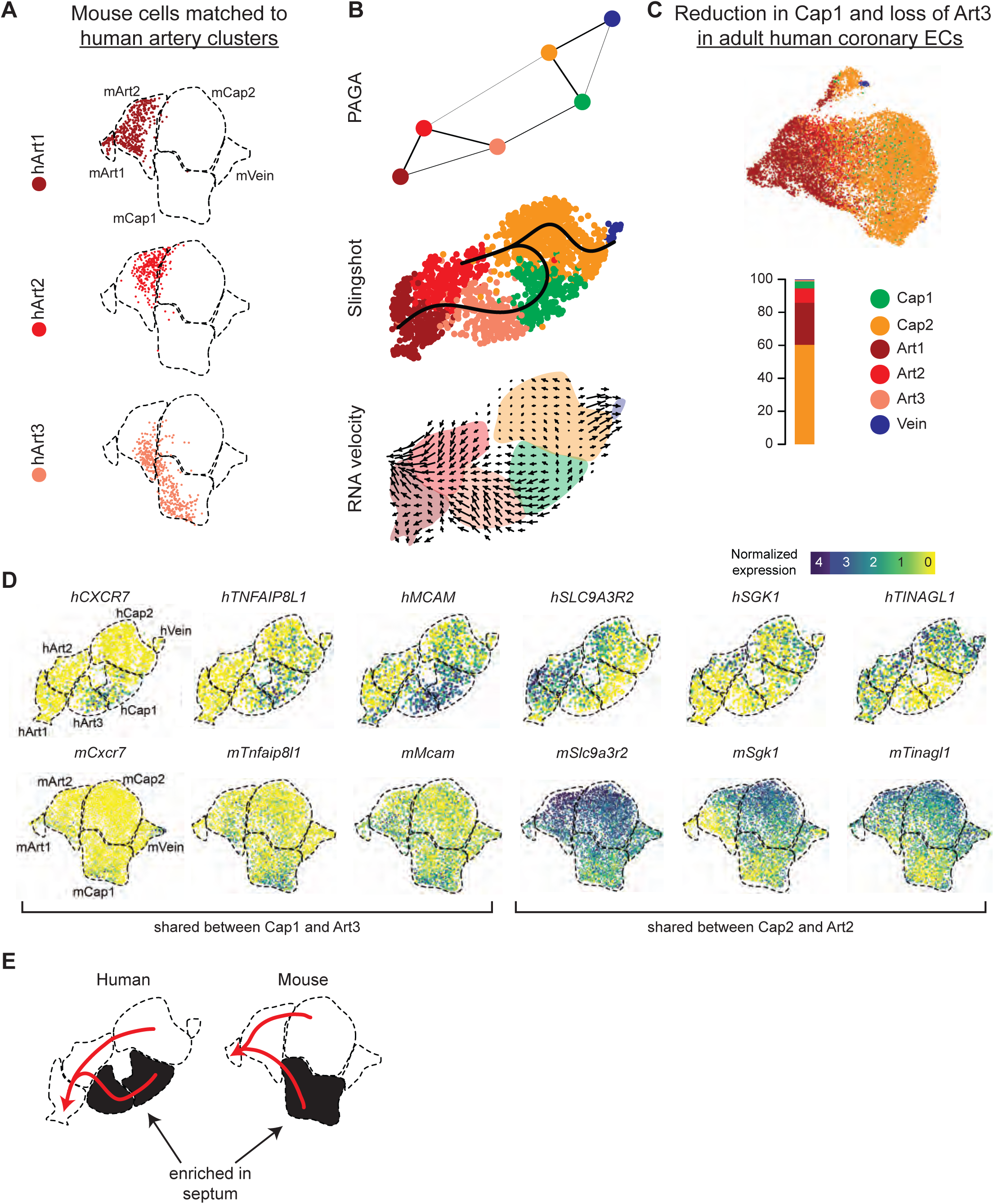
Trajectory analysis of developing human coronary arteries. (A) Reference mapping showed that e17.5 mouse ECs from Fig. 5g were assigned to all three human artery subsets. (B) Trajectory analysis of human coronary ECs using PAGA, Slingshot, and RNA velocity suggested that artery ECs are formed by capillary EC differentiation, as in mice. (C) Reference mapping adult human coronary ECs from a publicly available dataset to human fetal ECs showed that most mature cells match to Art1, Art2 or Cap2. (D) UMAPs showing expression of selected genes shared between hCap1 and hArt3 and hCap2 and hArt2, in both human and mouse. Previously defined clusters are outlined. (E) Schematic illustrating enrichment of septum ECs in mouse Cap1 and human Cap1 and Art3, as well as trajectories from capillary to artery in both human and mouse.

Reference mapping from the fetal to adult human datasets also revealed a notable reduction in Cap1 cells (28% of fetal capillary cells are Cap1 versus 6% of adult capillary cells)(Fig. 6c). In mice, e17.5 Cap1 and Cap2 converged into a relatively homogenous capillary population in adult (Fig. 1i). The small percentage of adult human coronary ECs mapping to the fetal Cap1 cluster, as well as the spatial overlap between cells mapping to Cap1 and Cap2 in the adult UMAP reduction, indicates that these developmental states related to flow and oxygenation also converge in humans.

We next performed trajectory analysis to investigate whether arteries develop through the differentiation of capillary ECs, which is the developmental pathway in mice (Red-Horse et al., 2010; Su et al., 2018). Two common methods for estimating trajectories, PAGA and Slingshot, identified connections between human Cap1 and Art3 and Cap2 and Art2 (Fig. 6b and S11b). RNA velocity suggested directionality going from the capillaries into arteries (Fig. 6b). These two predicted capillary to artery transitions suggested that Cap1/Art3 and Cap2/Art2 may be differentiation trajectories occurring in two different locations in the heart, i.e. septum vs. ventricle walls. This is supported by the observation that several genes are specifically co-expressed in Cap1 and Art3 (including *CXCR7*, *MCAM*, and *TNFAIP8L1*, *PGF)* or in Cap2 and Art2 (including *TINAGL1*, *SLC9A3R2*, *SGK1*, *THBD*, *LIMS2*, *CALCRL),* some of which were shared with mouse (Fig. 5h, 6d, and S10g). From these data, we propose a model where, in both mouse and human, two distinct subtypes of capillary cells at different locations in the developing heart initially produce two subtypes of artery cells, one of which eventually matures into the other (Fig. 6e).

### Characterization of human artery EC subpopulations

Since coronary artery disease is a leading cause of death and developmental information could suggest regenerative pathways, we next focused on the gene pathways present in developing human coronary arteries. As described above, unbiased clustering resulted in three artery states, each expressing known artery markers such as *GJA4* and *HEY1*, but also containing unique genes (Fig. 7a). The *SCENIC* package (Aibar et al., 2017), which uses gene expression information to identify transcription factor “regulons” present in cells, implicated *SOX17* as being strongly enriched in developing artery ECs (Fig. 7b), which is consistent with previous reports on artery development (Corada et al., 2013; Gonzalez-Hernandez et al., 2020). Transcription factors of potential importance that have not been previously implicated in artery development were *PRDM16* and *GATA2* (Fig. 7b). Interestingly, the *IRF6* regulon was specific to the most mature population suggesting a potential role in artery maturation (Fig. 7c). All of these regulons were similarly enriched in mouse artery cells (Fig. 7b and c). We also identified several genes with strong expression patterns in human artery ECs that were not found in mouse (Fig. 7d and Table 4). Interestingly, these included a GABA receptor, *GABBR2*, which was enriched in Art2, and a Glutamate receptor, *GRIA2*, which was is enriched in Art1. The human cells expressing *GABBR2* also co-expressed *SLC6A6*, a transporter that imports the GABBR2 ligand into cells (Tomi et al., 2008)(Fig. 7d). Finally, we localized different types of arteries in sections of human fetal hearts. In order to identify the artery subtypes, we used *in situ* hybridization for *GJA4* and *GJA5*. We found that *GJA5*-positive ECs, marking Art1, are in a small number of large arteries always covered with smooth muscle, while *GJA5*-negative/*GJA4*-positive ECs, marking Art2 and Art3, are numerous and in some cases not covered with smooth muscle (Fig. 7e and f). This observation is consistent with the trajectory showed in Fig. 6b, with the interpretation that the more mature *GJA5*+ arteries in human are larger and more proximal than *GJA5*- arteries, as they are in mice. This supports the conclusion that Art1 represents artery ECs in larger, more proximal branches, and Art2 and Art3 are smaller arterioles (Fig. 7g).

**Figure 7:**
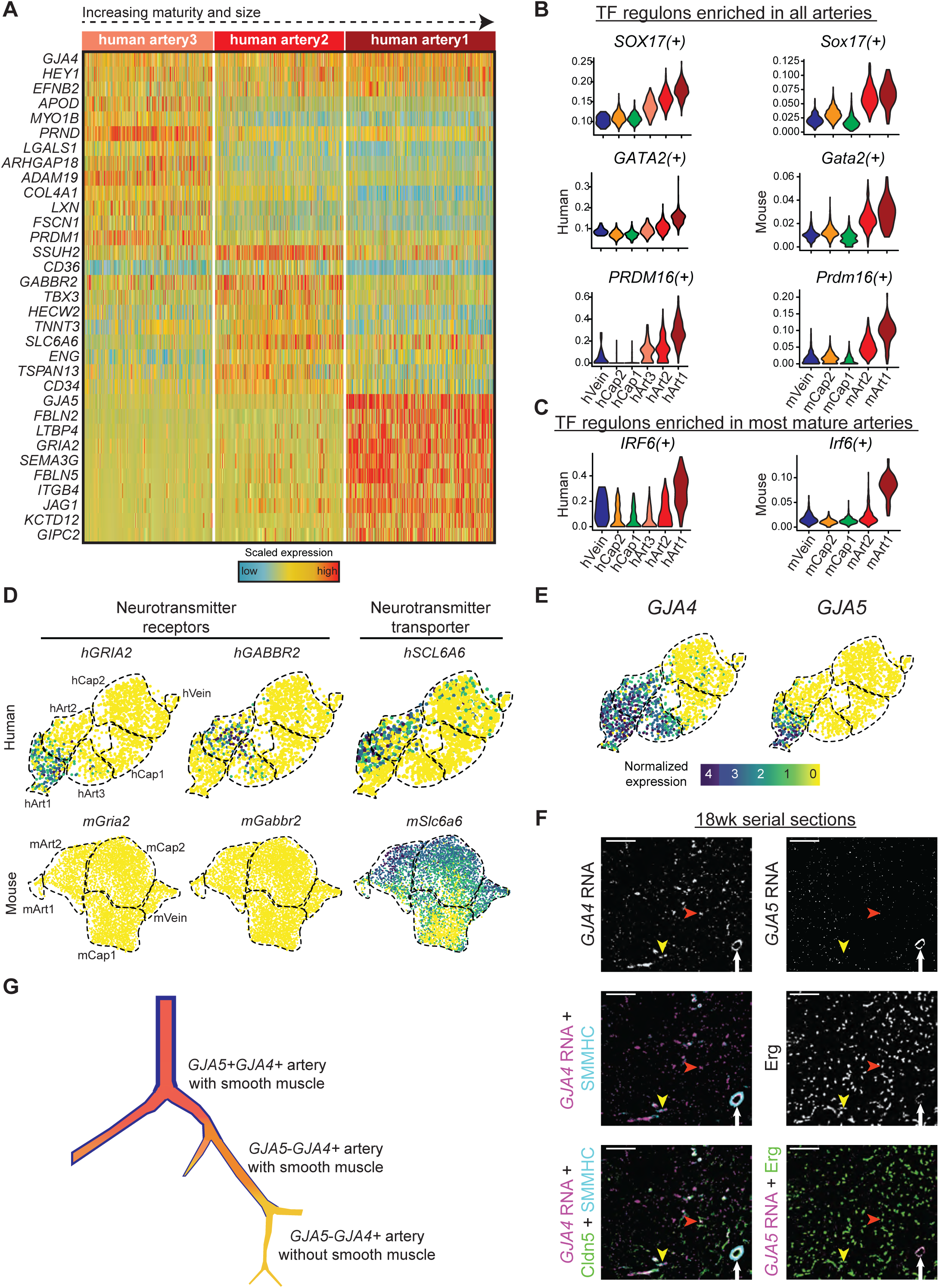
Gene expression in developing human coronary arteries. (A) Heatmap showing expression of selected genes enriched in human artery clusters. (B and C) Regulon scores from *SCENIC* analysis for TFs enriched in all human and mouse artery clusters (B) and for TFs enriched in human and mouse Art1 (C). (D) Human, but not mouse, developing coronary arteries expressed neurotransmitter receptors and their transporter. (E) *GJA4* and *GJA5* expression in human coronary ECs. (F) Serial sections from 18wk human fetal heart showing *in-situ* hybridization for the indicated mRNAs with immunofluorescence for the indicated proteins. Cluster hArt1 (*GJA5+GJA4+*) localizes to the largest arteries that are covered by mature SMMHC- positive smooth muscle (white arrows). Clusters hArt2 and 3 (*GJA4-GJA4*+) are smaller and can be either covered (yellow arrowheads) or not (red arrowheads) by smooth muscle. Scale bar = 100 μm. Scale bar from (E) also applies to (D). (G) Schematic of human coronary artery hierarchy.

**Table 4.**
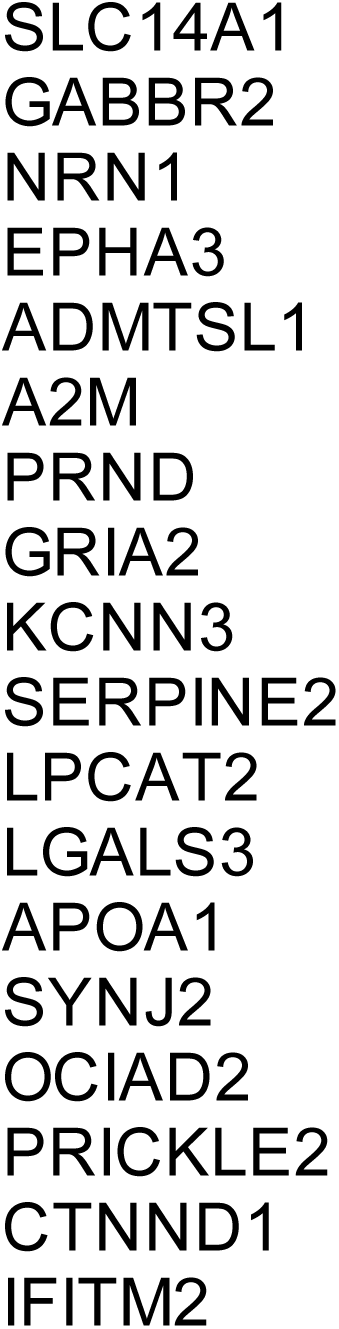
Genes unique to human coronary ECs.

## Discussion

Coronary ECs differentiate predominantly from the Endo and the SV. Although the spatial arrangement of these two lineages in the developing and adult mouse heart have been well characterized, it was previously unknown if either the distinct origin or localization of Endo- and SV-derived cells results in transcriptional or functional differences. Here, we used scRNAseq of lineage-traced ECs to address this question. At e12, when coronary vessels are just beginning to form, ECs transcriptionally segregated into two groups which are correlated with either Endo- or SV-specific expression patterns, and one of which is composed of only SV-lineage cells. However, at e17.5, during a phase of rapid growth and vascular remodeling, ECs segregated primarily based on being localized to either the septum or ventricular walls, the former of which expressed a genetic signature of experiencing low oxygen and blood flow. There was also differential gene expression between the dorsal and ventral walls of the developing heart. This could be due to differences in the timing and degree of their vascularization. We previously showed that coronary vessels are more numerous and provide more coverage on the dorsal side of the heart compared to the ventral side, at least until e15.5 (Red-Horse et al., 2010; Chen et al., 2014). Therefore, whatever environmental variables related to blood supply (including flow and hypoxia) distinguish the septum and the dorsal wall at e17.5 likely also cause the differences between the ventral and dorsal walls show in Fig 3d-f. In adult hearts, Endo- and SV-derived ECs cannot be distinguished either by gene expression or by their level of proliferation in response to IR injury. Altogether, these findings demonstrate that over the course of embryonic and post-natal development, coronary ECs from separate lineages converge both transcriptionally and functionally.

This result is relevant to future studies aiming to use developmental pathways to enhance regeneration in adult hearts. For example, it implies that approaches to stimulate re-growth of the vasculature after myocardial injury will affect all cells equally with regard to lineage, and that achieving vascular remodeling in adults may require replicating specific environmental cues and signals (especially those related to hypoxia and flow). The transcriptional similarity of adult Endo- and SV-derived cells helps explain the prior observation that mutant embryos whose coronary vasculature was derived primarily from the Endo due to compensation for loss of SV sprouting grew into phenotypically normal adults despite developmental defects (Sharma et al., 2017). It also raises additional questions for investigation, namely, how is it that coronary ECs can over time lose the signatures both of their progenitors (between e12 and e17.5) and of their developmental “home” (between e17.5 and adult)? Additional approaches, including ATAC-seq, could aid in addressing this question. The unexpected plasticity of these embryonic cells in their ability to become completely transcriptionally identical represents a potential model for efforts to stimulate faithful differentiation of very specific cell types from induced pluripotent stem cells or non-canonical progenitors (Yamanaka, 2020).

Another outstanding question addressed by this study is the degree of transcriptional similarity between developing mouse and human coronary ECs. Several groups have recently used scRNAseq to profile cell types in the fetal human heart, and identified individual genes that are enriched in either mouse or human (Asp et al., 2019; Cui et al., 2019; Miao et al., 2020; Suryawanshi et al., 2020). Here, our experiments enriched for ECs to enable a high-resolution comparison of this cellular compartment. The data showed that mouse transcriptional clusters specifically enriched in either septal or free wall cells are reproduced in our 11, 14, and 22 week human scRNAseq datasets, and many of the defining genes are conserved. Unlike in mouse, there is no apparent difference between ventral and dorsal gene expression patterns, indicating that the timing of vascularization might be more consistent throughout the developing human heart. Furthermore, the observation that adult human capillary ECs almost completely map to developmental Cap2 (Fig. 6c) indicates that the hypoxic and low flow Cap1 is also resolved in adult human hearts as shown in mice. Although it is not possible to definitively identify the origins of human heart ECs using scRNAseq alone, the presence of similar cell states between mouse and human at these timepoints, as well as the convergence of adult capillary cells in both, lends confidence to the notion that coronary development generally follows the same progression in these two species.

Despite overall similarities in cell types, the scRNAseq analysis did reveal some interesting differences in gene expression between mouse and human, some of which may explain the anatomical differences in their vasculature. For instance, human Art1 and Art2 specifically expressed the glutamate receptor *GRIA2* and the GABA receptor *GABBR2*, respectively. It was previously demonstrated that exposure of ECs to GABA *in vitro* led to a reduction in response to inflammatory stimulus (Sen et al., 2016), and that mutations in brain ECs of a different GABA receptor, *Gabrb3*, resulted in defects in neuronal development *in vivo* (Li et al., 2018). These genes were not present in human adult coronary artery ECs. Further investigation may reveal an important role for GABA and glutamate signaling in human coronary development.

In summary, we have shown that in both mouse and human, phenotypically distinct lineage- and location-based cell states of coronary ECs converge in adults, and that embryonic lineage does not influence injury responses. This is a demonstration of the significant plasticity of the vasculature and the influence environmental factors have in shaping heterogeneity, as well as the strength of mice as a model organism for human heart development.

## Methods

### Mice

#### Mouse strains

All mouse husbandry and experiments were conducted in compliance with Stanford University Institution Animal Care and Use Committee guidelines. Mouse lines used in this study are: *BmxCreER* (Ehling et al., 2013), *ApjCreER* (Chen et al., 2014), *mTmG* (The Jackson Laboratory, *Gt(ROSA)26Sortm4(ACTB-tdTomato*,*-EGFP)Luo*/J, Stock #007576), *tdTomato* (The Jackson Laboratory, B6.Cg-Gt(ROSA)26Sortm9(CAG-*tdTomato*)Hze/J, Stock #007909), and CD1 (Charles River Laboratories, strain code: 022).

#### Breeding and tamoxifen administration

Timed pregnancies were determined by defining the day on which a plug was found as e0.5. For Cre inductions, Tamoxifen (Sigma-Aldrich, T5648) was dissolved in corn oil at a concentration of 20 mg/ml and 4 mg was administered to pregnant dams using the oral gavage method on days e8.5 and e9.5 (Fig. 1a, Fig. 3d-e, Fig. S1a) or day e11.5 (Fig. 1b). Combined injections of tamoxifen at e8.5 and e9.5 led to labeling of 94.44% of Endo cells and 3.61% of SV cells at e12.5. For the adult injury experiments, either 4 mg of Tamoxifen or 1 mg of 4-OH Tamoxifen (Sigma-Aldrich, H6278) was delivered on day e9.5 or e10.5 (Fig. 4a, Fig. S1b), respectively. The five *BmxCreER*-*Rosa^tdTomato^* mice used for the adult 10X experiment were 13 weeks old and all male. The 11 *BmxCreER*-*Rosa^tdTomato^* adult mice used for the injury experiments were 12 weeks old and all male (Fig. 4a). The three CD1 adult mice used for quantification of EC proliferation in uninjured hearts were 10 weeks old and all female (Fig. 4e,f,h). The three CD1 adult mice used for Car4 staining were 6 weeks old and all female (Fig. 3h). Adult mice for the scRNAseq and injury experiments were obtained by harvesting litters at e18.5 and fostering pups with a different female who had given birth 0-4 days earlier.

### Human hearts

Under IRB approved protocols, human fetal hearts were collected for developmental analysis from elective terminations. Gestational age was determined by standard dating criteria by last menstrual period and ultrasound (Obstetrics, 2009). Tissue was processed within 1h following procedure. Tissue was extensively rinsed with cold, sterile PBS, and placed on ice in cold, sterile PBS before further processing as described below. Pregnancies complicated by multiple gestations and known fetal or chromosomal anomalies were excluded.

### Single-cell RNA sequencing protocol

#### e12, e17.5, and adult mouse scRNAseq

*BmxCreER*-*Rosa^tdTomato/tdTomato^* males were crossed to CD1 females, which were dosed with tamoxifen at e8.5 and e9.5 (e12 and e17.5) or with 4-OH tamoxifen at e10.5 (adult). Either early in the day on e12, or midday on e17.5, embryos were removed and placed in cold, sterile PBS. 42 Cre+ e12 embryos and 9 Cre+ e17.5 embryos were identified by their fluorescent signal and used for single-cell isolation. 5 Cre+ adult males were identified by Cre amplification and used for single-cell isolation. Hearts were isolated and dissected to remove the atria and outflow tract, keeping the ventricles, SV, and valves (e12 and e17.5) or to remove the atria, outflow tract and valves, keeping the ventricles (adult). Hearts were then dissociated in a 600 μl mix consisting of 500 U/ml collagenase IV (Worthington #LS004186), 1.2 U/ml dispase (Worthington #LS02100), 32 U/ml DNase I (Worthington #LS002007), and sterile DPBS with Mg2+ and Ca2+ at 37 degrees for 45 minutes and resuspended by pipetting every 5 minutes. Once digestion was complete, 5 mL of a cold 5% FBS in PBS mixture was added and the suspension was filtered through a 40 micron strainer. After further rinsing the strainer with 5 mL of 5% FBS/PBS, the cell suspension was centrifuged at 400*g* at 4°C for 5 min. The cells were washed and resuspended once more in 1 mL 5% FBS/PBS. The following antibodies were added at the concentration of 1:50 and incubated on ice for 45 minutes: APC/Cy7 Cd45 (Biolegend #103116), APC Pecam1 (Biolegend #102410), APC/Cy7 Ter-119 (Biolegend #116223). DAPI (1.1 μM) was added to the cells immediately before FACS. Once stained, the cells were sorted on a Aria II SORP machine into 1.5 mL tubes. The gates were set up to sort cells with low DAPI, low Cd45 (hematopoietic cells), low Ter119 (erythroid cells), high Pecam1 (endothelial marker), and either high or low PE-Texas Red (*tdTomato* positive or negative). Compensation controls were set up for each single channel (PE-Texas Red, APC, APC/Cy7) before sorting the final cells. The samples were then submitted to the Stanford Genome Sequencing Service Center for 10x single-cell v3.1 3’ library preparation. For each stage, libraries from the *tdTomato* positive and negative samples were pooled and sequencing was performed on two lanes of a Illumina NovaSeq 6000 SP flow cell.

#### 22 week fetal human heart scRNAseq

The experiment was performed using the same procedure as the mouse samples unless noted here. The heart was kept in cold, sterile PBS. It was dissected to remove the atria, outflow tract, and valves, keeping only the ventricles. Dissociation was performed as described for the mouse samples except that multiple tubes of the 600 μl mix were used for each heart. The antibodies used for staining were: Pacific Blue CD235a (Biolegend #349107), FITC CD36 (Biolegend #336204), APC/Cy7 PECAM1 (Biolegend #303119), Pacific Blue CD45 (Biolegend #304021). The gates were set up to sort cells with low DAPI, low CD45 (hematopoietic cells), low CD235A (erythroid cells), high PECAM1 (endothelial marker), and low FITC. After staining each cell was sorted into a separate well of a 96-well plate containing 4 μl lysis buffer. Cells were spun down after sorting and stored at −80 °C until cDNA synthesis.A total of 1920 PECAM1+CD36- and PECAM1+ cells were sorted and processed for cDNA synthesis. Cells were analyzed on the AATI 96-capillary fragment analyzer, and a total of 1382 cells that had sufficient cDNA concentration were barcoded and pooled for sequencing.

#### 11 and 14 week fetal heart scRNAseq

The experiment was performed using the same procedure as the 22 week heart unless noted here. The antibodies used for staining were: PerCP/Cy5.5 CD235a (Biolegend #349110), PerCP/Cy5.5 CD45 (Biolegend #304028), FITC CD36 (Biolegend #336204), APC/Cy7 PECAM1 (Biolegend #303119). The gates were set up to sort cells with low DAPI, low CD45 (hematopoietic cells), low CD235A (erythroid cells), high PECAM1 (endothelial marker), and either low or high FITC. A total of 1824 PECAM1+CD36-, PECAM1+CD36+ and PECAM1+ cells from the 11 week heart and 1920 PECAM1+CD36-, PECAM1+CD36+ and PECAM1+ cells from the 14 week heart were sorted and processed for cDNA synthesis. A total of 1530 11 week and 1272 14 week cells that had sufficient cDNA concentration were barcoded and pooled for sequencing.

Synthesis of cDNA and library preparation for the fetal human heart cells was performed using the Smart-seq2 method as previously described (Picelli et al., 2014; Su et al., 2018). Libraries from the fetal human heart cells were part of a pool of samples that was sequenced on four lanes of a Illumina NovaSeq 6000 S4 flow cell.

### Single-cell RNA sequencing data analysis

#### Processing of sequencing data

Raw Illumina reads for all datasets were demultiplexed and converted to FASTQ using *bcl2fastq* (Illumina). For human, sequencing adapter and PCR primer sequences were trimmed from reads using cutadapt 2.7 (Martin, 2011). For mouse, reads were aligned to GRCm38 Ensembl release 81 as well as *EGFP* and *tdTomato* sequences and a gene count matrix was obtained using Cell Ranger v3.1.0 (10X Genomics). For human, reads were aligned with STAR v2.7.1a (Dobin et al., 2013) to GRCh38 Ensembl release 98, and a gene count matrix was obtained using the *featureCounts* function of Subread v1.6.0 (Liao et al., 2014).

#### Processing of count data

The majority of scRNAseq data analysis was performed using R and Seurat v3 (Stuart et al., 2019). Cells were deemed low-quality and excluded from downstream analysis if they expressed less than 1000 genes or if more than 10% of reads aligned to mitochondrial genes. A small number of cells were removed from the mouse e17.5 and adult *tdTomato* negative samples which were expressing *tdTomato*. For all datasets, non-endothelial subtypes (e.g. blood and immune cells, cardiomyocytes, smooth muscle, fibroblasts) as well as a small number of lymphatic cells were removed. For the adult mouse, endocardial cells were removed, as well as a small cluster of cells enriched in dissociation-induced genes (ex. *Hspa1a, Hspa1b, Socs3, Junb, Atf3*) (van den Brink et al., 2017). To obtain the subsets of vascular ECs shown in Fig. 1, SV, valve, endocardium, SV, and cycling cells were removed as shown in Fig. S2b. Additionally, a cluster of cells with a lower gene count and higher mitochondrial percentage were removed from the e17.5 dataset. The cells used for the cell cycle analysis in Fig. S4 include all the cells used Fig. 1f combined with the non-endocardial cycling cells shown in Fig. S2b.

Count data from Su et al (Su et al., 2018), and Litvinukova et al (Litvinukova et al., 2020) were used to analyze gene expression in e12.5 mouse, and adult human hearts, respectively. From these datasets only vascular endothelial cells were retained, excluding endocardium, SV, valve endothelium, lymphatic endothelium, and non-endothelial cell types.

Normalization, variable feature selection, scaling, and dimensionality reduction using principal component analysis were performed using the standard Seurat v3 pipeline (Stuart et al., 2019). For the e17.5 and adult mouse datasets the technical variables genes per cell, reads per cell, and mitochondrial read percentage were regressed out in the *ScaleData* function. Following this, construction of a shared nearest neighbor graph, cluster identification with the Louvain algorithm (Stuart et al., 2019), and Uniform Manifold Approximation and Projection (UMAP) dimensionality reduction (Becht et al., 2018) were performed using the *FindNeighbors*, *FindClusters*, and *RunUMAP* functions in Seurat using the parameters listed below. Clustering resolution was determined individually for each dataset as the highest resolution at which every cluster expressed at least one unique marker gene:

Fig. S2a (11520 cells)- 25 dimensions, Louvain resolution = 0.8
Fig. S2b (12,205 cells)- 25 dimensions, Louvain resolution = 0.8
Fig. 1c (436 cells)- 25 dimensions, Louvain resolution = 0.6
Fig. 1f (4801 cells)- 25 dimensions, Louvain resolution = 0.4
Fig. 1i (649 cells)- 25 dimensions, Louvain resolution = 0.3
Fig. S7a (356 cells)- 20 dimensions, Louvain resolution = 1
Fig. S4a (8495 cells)- 25 dimensions, Louvain resolution = 0.4 (after regression), Louvain resolution = 0.6 (before regression)
Fig. 5b (2339 cells)- 30 dimensions, Louvain resolution= 1.4
Fig. 5c (1586 cells)- 20 dimensions, Louvain resolution = 1
Fig. 6c (13,476 cells)- 50 dimensions, Louvain resolution = 0.8

For the human dataset, clustering with a resolution of 1 resulted in 7 clusters: Art 1-3, Veins, Cap1, and two additional clusters which were eventually merged into Cap2. The reason for this merging is that one of these clusters was composed only of cells from the 22 week sample, and one of the top differentially expressed genes between these two clusters was *XIST* (as the 22 week sample was the only one expressing *XIST*, we concluded it was the only male sample). In addition, most of the other genes distinguishing these two clusters were dissociation-induced genes as described by van den Brink et al (van den Brink et al., 2017). Thus, it was determined that the difference between these two clusters was not biologically meaningful (mainly dissociation effects which were different between samples), and they were treated as one.

#### Differential expression testing

Differential gene expression testing was performed with the *FindMarkers* and *FindAllMarkers* functions in Seurat using the Wilcoxon Rank Sum test. All differential genes were defined using parameters logfc.threshold = 0.3, min.pct = 0.2 and filtered for p-value < 0.001.

#### Cell cycle regression

Cell cycle regression for Fig. S4 was performed using top 100 gene markers for the cycling clusters by p-value based on Wilcoxon Rank Sum test and the vars.to.regress parameter in the Seurat *ScaleData* function.

#### Pearson correlation

Pearson correlation heatmaps in Fig. 2e-f were created with the *heatmaply_cor* function from heatmaply (Galili et al., 2018).

#### Dataset reference mapping

For cross-dataset mapping in Fig. 5, the *getLDS* function in biomaRt (Durinck et al., 2005; Durinck et al., 2009) was used to identify every human gene that has a corresponding mouse gene and vice versa. Genes were only retained if they had a 1:1 mapping between human and mouse. The raw counts matrix for the human fetal data (all three sages pooled) was then filtered to include only these genes, and only the cells used in Fig. 5c, and a new Seurat object was created from this count matrix. Similarly the raw count matrix for the e17.5 mouse data was filtered to include only these genes, and only the cells used in Fig. 1f, and a new Seurat object was created from this count matrix. Standard normalization and scaling was performed in Seurat. To perform the mappings between datasets in Figs. 5e-g, 6c, and S10f, the Seurat functions *FindTransferAnchors* and *TransferData* were performed using 30 dimensions and Canonical Correlation Analysis dimensionality reduction. The label transfer method employed is described in detail by Stuart et al (Stuart et al., 2019). Briefly, diagonalized canonical correlation analysis is used for dimensionality reduction of both datasets, followed by L2 normalization. Then, a mutual nearest neighbors approach is used to identify pairs of cells (“anchors”) between the two datasets that represent a similar biological state. Every cell in the query dataset is assigned an anchor in the reference dataset (with an associated anchor score), and the cluster label of the reference cell is assigned to the query cell.

The major limitation of this method as we applied it is that it is being used to compare datasets from different species that were generated using different methods (Smart-seq2 and 10x) and different sequencing depths. These differences as well as limitations associated with these methods impact the accuracy of the label transfer. For example, most of the mouse vein cells match to human Cap2 rather than human veins (Fig 5g), likely because there are so few human vein cells while cells in the larger Cap2 population have a larger local neighborhood to strengthen their anchor score. In addition, the matching is limited to the clusters that are pre-annotated in each dataset (i.e., each human cell will match to its closest mouse cluster, even if there is not a true biological correlate). Finally, genes used in the mapping were limited to those that had a 1:1 homology mapping between mouse and human, meaning that some information was eliminated before label transfer.

#### Trajectory analysis

Trajectory analyses shown in Figs. 6b and S11b and were performed with PAGA (Wolf et al., 2019) (filtered for edge weight greater than or equal to 0.14), RNA velocity (La Manno et al., 2018) (using the python function *veloctyto run-smartseq2*, followed by the R package velocyto.R with the function *show.velocity.on.embedding.cor* with fit.quantile = 0.05, grid.n = 20, scale = ‘sqrt’, arrow.scale = 3, and n= 50-100), and Slingshot (Street et al., 2018) (using the *slingshot* function with Cap2 as the starting cluster and stretch = 1)

#### Transcription factor enrichment

Transcription factor enrichment was performed with SCENIC (Aibar et al., 2017) using the *pyscenic* functions and the recommended pipeline (Van de Sande et al., 2020). The loom file output from SCENIC was then imported into Seurat, and the *FindAllMarkers* function with the Wilcoxon Rank Sum test was used to identify differential regulons between clusters.

### Immunofluorescence and imaging

#### Tissue processing and antibody staining

E17.5 embryos were dissected in cold 1X PBS and fixed in 4% PFA for 1 hour at 4°C, followed by three 15 minute washes in PBS. Hearts were then dissected from the embryos. Adult mouse hearts were dissected and fixed in 4% PFA for 4 -5 hours at 4°C, followed by three 15 minute washes in PBS. Hearts were dehydrated in 30% sucrose overnight at 4°C, transferred to OCT for a 1 hour incubation period, and frozen at -80°C. For each heart, the whole ventricle was cut into 20 μm thick sections which were captured on glass slides. Staining was performed by adding primary antibodies diluted in .5% PBT (.5% Triton X-100 in PBS) with .5% donkey serum to the sections and incubating overnight at 4°C. The following day the slides were washed in PBS 3 times for 10 minutes followed by a 2 hour room temperature incubation with secondary antibodies, three more 10 minute washes, and mounting with Fluoromount-G (SouthernBiotech #0100-01) and a coverslip fastened using nail polish. Human fetal hearts were fixed in 4% PFA for 24-48 hrs at 4°C, followed by three 15 minute washes in PBS. The hearts were sequentially dehydrated in 30%, 50%, 70%, 80%, 90% and 100% ethanol, washed three times for 30 minutes in xylene, washed several times in paraffin, and finally embedded in paraffin which was allowed to harden into a block. For each heart, the whole ventricle was cut into 10 μm thick sections which were captured on glass slides.

#### In situ hybridization for GJA4 and GJA5

RNA was isolated from a 23 week human fetal heart using Trizol-based dissociation followed by the RNEasy Mini Kit (Qiagen #74104). cDNA was created from this RNA using the iScript Reverse Transcription Supermix (Bio-Rad #1708840). Primers used to amplify *GJA4* are: 5’-AAACTCGAGAAGATCTCGGTGGCAGAAGA-3’ and 5’- AAATCTAGACTGGAGAGGAAGCCGTAGTG-3’. Primers used to amplify *GJA5* are: 5’- AAACTCGAGAATCAGTGCCTGGAGAATGG-3’ and 5’-AAATCTAGATGGTCCATGGAGACAACAGA-3’. Digoxin-linked probes were transcribed using the Roche DIG RNA Labeling Kit (Millipore Sigma #11175025910). In-situ hybridization was performed as previously described (Koop et al., 1996) with a modification to develop the fluorescent signal. Briefly, after hybridization, sections were incubated overnight at 4°C with anti-DIG POD (Millipore Sigma #11207733910). The next day, sections were washed 4x 1 hour in 1X MABT. Finally, sections were washed for 3x 10 minutes with 0.1M Borate buffer pH 8.5 and stained with bench-made tyramide (Vize et al., 2009).

*In situ hybridization for TINAGL1*. *In situ* hybridization was performed using the RNAscope Multiplex Fluorescent V2 assay (Advanced Cell Diagnostics #323100), with probe for human *TINAGL1*(Advanced Cell Diagnostics #857221-C2) and OPAL 570 fluorophore (Akoya #FP1488001KT), following the manufacturer’s protocol.

#### Microscopy and Image Processing

Images were captured on a Zeiss LSM-700 confocal microscope. For each experiment, littermate embryos were stained together and all samples were imaged using the same laser settings. For each experiment, laser intensity was set to capture the dynamic range of the signal. Images were captured using Zen (Carl Zeiss) and processed using FIJI (NIH) and Illustrator (Adobe). Any changes to brightness and contrast were applied equally across the entire image. All mouse imaging experiments were performed with at least three individual samples.

#### Antibodies

The following primary antibodies were used: anti-ERG (1:200; Abcam, ab92513), anti-Car4 (1:200; R&D, AF2414), anti-Smmhc (1:100; Proteintech, 21404-1-AP), anti-Cldn5 (1:100; Invitrogen, 35-2500). Secondary antibodies were Alexa fluor-conjugated antibodies (488, 555, 633) from Life Technologies used at 1:250.

#### Quantification

Quantification of Car4 (Fig. 3d-e, Table 3) and EdU (Fig. 4e-h) was performed using the CellCounter plugin in FIJI. For e17.5 embryos, Car4, Erg, and *tdTomato* were quantified in five ROIs in each of three sections from each of three hearts. The ROIs were 510 μm x 190 μm and were positioned to maximize coverage of the septum, left ventricular wall, right ventricular wall, dorsal wall, and ventral wall. For each heart, counts were combined across the three sections. For adult hearts, Car4 and Erg were quantified in two ROIs of 600 μm x 600 μm, one in the septum and one in the left ventricle.

For the lineage comparison in adult injured hearts (Fig. 4c-f), EdU, Erg, and *tdTomato* were quantified in 2 ROIs for each of 11 hearts at the level of the stitch (quantification from 2 ROIs was averaged), and in 1 ROI for each of 7 hearts at the apex. ROIs were 600 μm x 600 μm and were chosen to be in the region of the section with the greatest density of EdU staining, and whenever possible, to span portions of both the inner and outer myocardial wall. For uninjured controls for the lineage comparison, one ROI of 600 μm x 600 μm was chosen in both the middle of the myocardial wall of the left ventricle, and at the apex, from each of three uninjured 6 week female CD1 mouse hearts. *BmxCreER*-*Rosa^tdTomato^* was observed to label ECs in large arteries in adult tissues even without tamoxifen administration (Red-Horse Lab, unpublished data). To compare proliferation in the inner and outer wall of the adult heart (Fig. 4f), one ROI each of 600 μm x 600 μm were chosen in a portion of the injured area (areas with high density of EdU staining, as indicated in Fig. 4b) overlapping with the inner or outer myocardial wall, in 3 injured hearts. In addition, one ROI each of 600 μm x 600 μm was chosen adjacent to the injured area, and overlapping with the inner or outer myocardial wall, in 3 injured hearts. Since the developmental origin of adult arteries cannot be determined using *BmxCreER*, large arteries were excluded from all ROIs during quantification for the lineage comparison after injury. However, we were able to use tdTomato labeling to identify arteries in the injury area for the quantification of artery proliferation. For each of 10 hearts at the level of the stitch, and for each of 4 hearts at the apex, large arteries labeled with tdTomato were identified in the injury area, and the number of Erg+ and Erg+Edu+ cells were counted. These were compared to counts of Erg+ and EdU+ cells in arteries identified in the left ventricles (3 arteries each) and apex (2 arteries each) of three uninjured 6 week female CD1 mouse hearts, with the arteries being identified by Smmhc staining.

For human *TINAGL1*, ROIs were captured at 40x magnification from one section each from an 18 week and a 20 week human fetal heart. For each section, 4 ROIs of 320 μm x 320 μm were chosen from the septum and left ventricle, and 3 ROIs of the same size were chosen from the right ventricle, ventral wall and dorsal wall. Automatic detection of Erg+ nuclei and probe spots were performed in QuPath using the default parameters with the following exceptions: cellExpansionMicrons=0.1, subcellular detection threshold=100. Measurements shown in Fig. 5k represent the values of “Num spots estimated”.

Graphs in Figs. 3 and 4, 5 and S8 were made in Prism 8.

#### Statistics

In Fig. 4e, paired t-tests were used to compare Endo-derived and SV-derived EC proliferation, and Welch’s t-test was used to compare Endo-derived or SV-derived EC proliferation with the uninjured control. Unpaired t-tests were used for Fig. 3e and 4f. Unpaired Welch’s t-tests were used for Fig. 4h. For Fig. 5k, comparisons were made using one-way ANOVA with Holm-Sidak’s multiple comparisons test.

### Ischemia reperfusion injury experiment

#### Surgery

*BmxCreER*-*Rosa^tdTomato/tdTomato^* males were crossed to CD1 females, which were dosed with tamoxifen e9.5 or with 4-OH tamoxifen at e10.5. In addition to being pharmacologically less toxic compared to Tam, 4-OHT is more potent at inducing Cre, given its stronger affinity for the ER domain in *CreER* strain (Robertson et al., 1982; Katzenellenbogen et al., 1984; Cardoso et al., 2003). By administering 4-OHT 1 day later than Tam, Cre was induced in all animals at approximately the same developmental time regardless of treatment. Pups were dissected from the pregnant dams at day e18.5 and fostered as described above. Ischemia-reperfusion (IR) was performed in 12-week old mice by the Stanford Murine Phenotyping Core (SMPC) that is directed by Dr. Dan Bernstein. To summarize, mice were anesthetized using isoflurane and placed on a rodent ventilator to maintain respiration before opening the chest cavity. The left anterior descending coronary artery (LAD) was ligated with a 8.0 silk suture and resulting ischemia of the myocardium was verified by blanching the left ventricular wall. After 40 min, the suture was removed around the LAD, allowing for the reperfusion of downstream myocardium. To end the procedure, the chest was closed, and post-operative analgesia was administered to the mice.

#### In vivo proliferation assay

To assess EC proliferation after IR, 5-ethynyl-2′-deoxyuridine (EdU) (Thermo Fisher Scientific, cat. #E10415) was diluted 2.5 mg/ mL in sterile PBS and injected intraperitoneally on days 3 and 4 post-injury at a dose of 10 μL/ g body weight. Hearts were harvested from sacrificed animals on day 5 post-injury. Cryosectioning and antibody staining for Erg was performed as described above. To detect endothelial proliferation, the protocol for Click-iT^®^ EdU Imaging (Thermo Fisher Scientific, cat. #C10086) was carried out according to manufacturer instructions.

## Acknowledgements

K.R. is supported by the NIH (R01-HL128503). R.P. is supported by an AHA graduate fellowship. Sequencing of the adult mouse and fetal human datasets was funded by the Chan Zuckerberg Biohub. We thank Ralf Adams for sharing the *BmxCreER* mouse line. We thank the Stanford Family Planning Clinic and Purnima Iyer Narasimhan for assistance with tissue procurement. We thank Gavin Sherlock for access to equipment needed for single-cell library preparation. We thank all members of the Red-Horse lab for technical and intellectual support. We thank Rahul Sinha, Anshul Kundaje and Laksshman Sundaram for discussion and advice about scRNAseq technique and analysis. We thank Biafra Ahanonu for discussion and advice about figure and manuscript preparation. We thank members of the Stanford Genome Sequencing Services Center which is supported by NIH Grant # 1S10OD020141-01. V.D.W. is supported by the H&H Evergreen Fund.

## Contributions

R.P., G.D., and K.R. conceptualized the study and wrote the manuscript. R.P. and G.D. performed experiments. R.P., J.K. and M.Z. performed cardiac injury study. R.P. performed computational analysis. S.S.K., R.C.J., S.R.Q., I.W., V.D.W., and D.B. assisted with experiments and provided experimental resources.

---

**Supplementary Figure 1:**
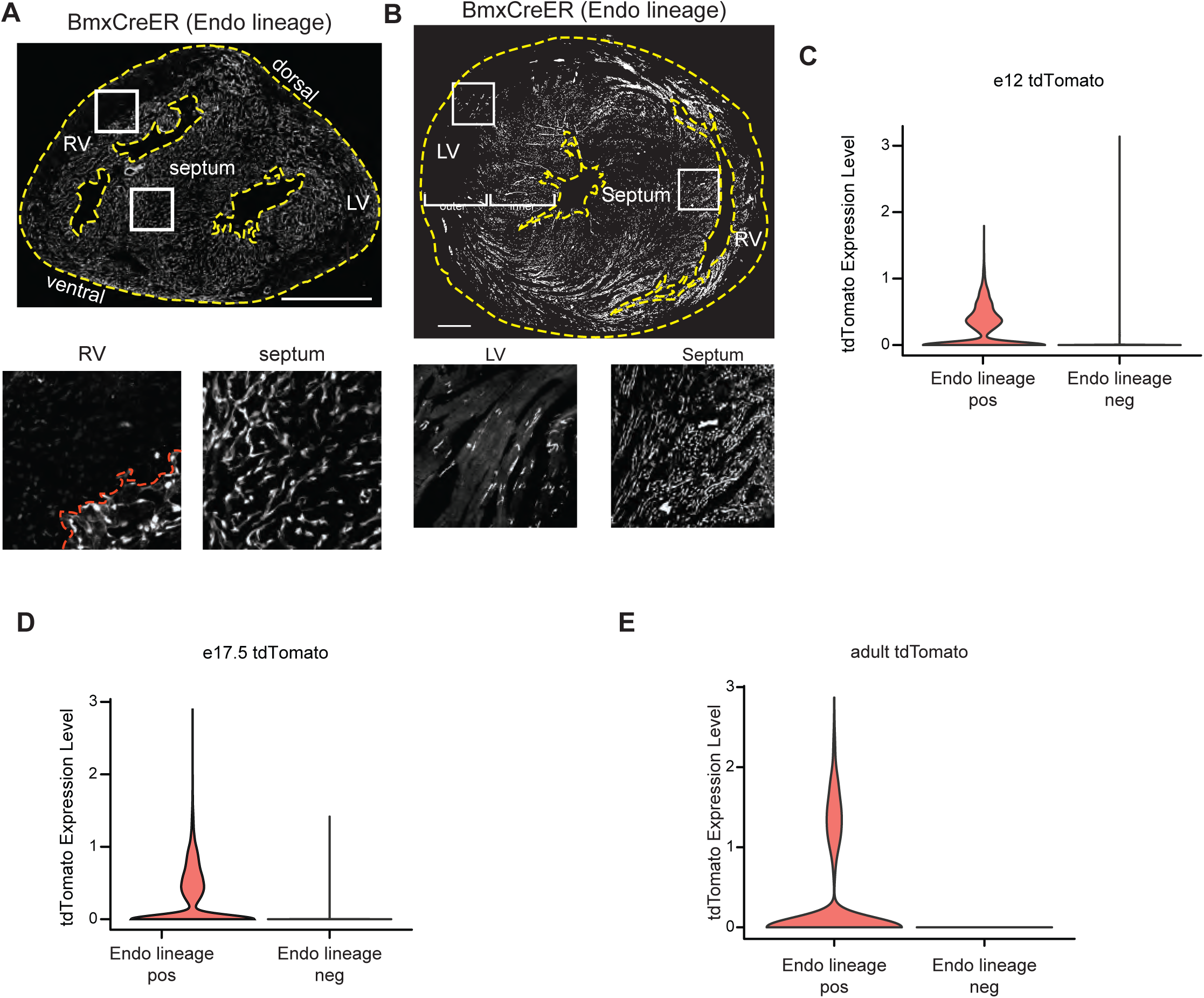
Localization and expression of recombinant markers. (A and B) *tdTomato* localization in sections *BmxCreER;Rosa^tdTomato^* hearts at e17.5 (A) and adult (B). Dashed yellow lines indicate the borders of the tissue sections. Dashed red line indicates the CV/Endo border. (C-E) Expression of the *tdTomato* gene in the Endo-enriched (Endo lineage positive) and SV-enriched (Endo lineage negative) sorted samples at e12 (C), e17.5 (D), and adult (E) from *BmxCreER;Rosa^tdTomato^* hearts (as shown in Fig. 1a-b). Scale bars = 500 μm.

**Supplementary Figure 2:**
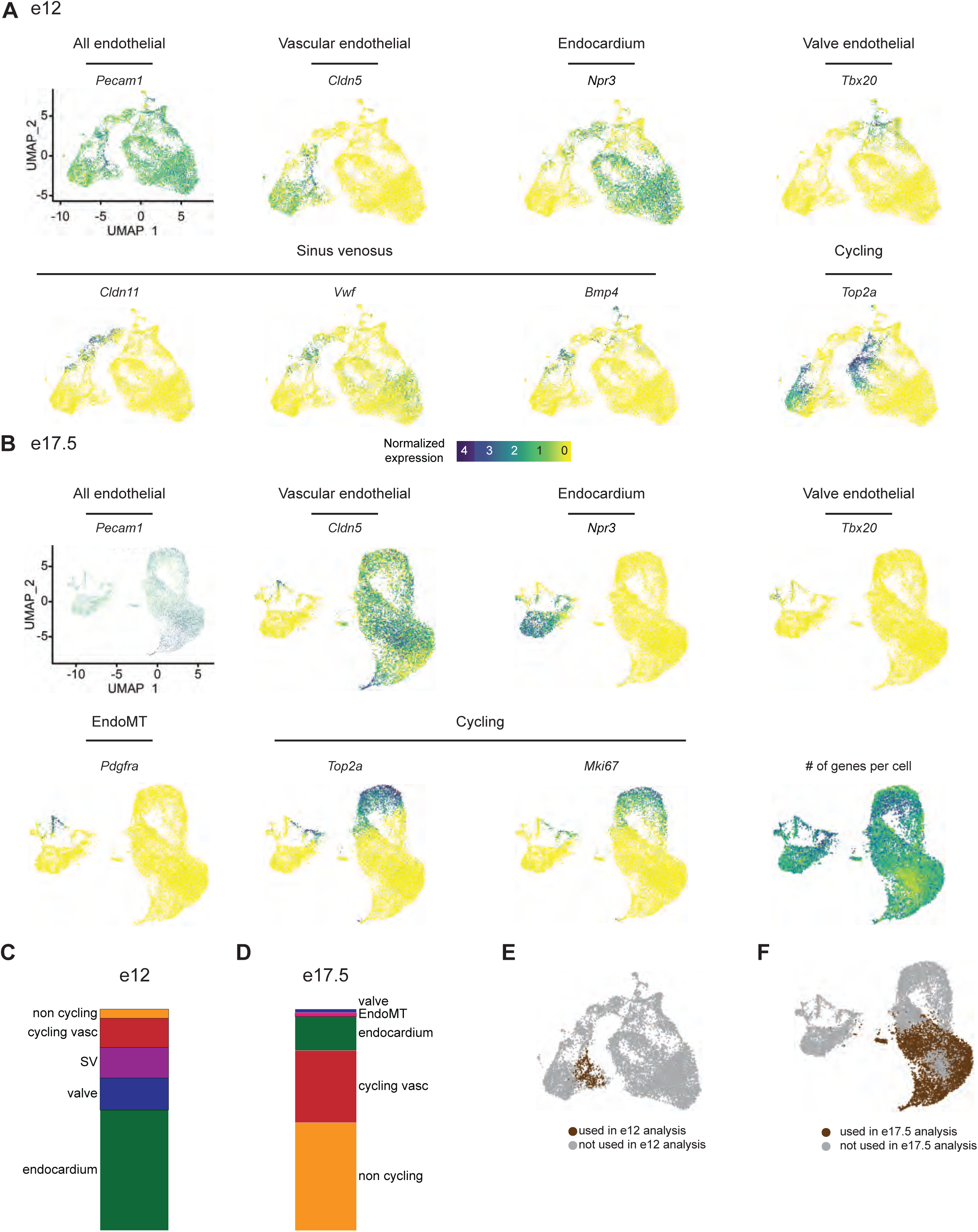
Selection of coronary vascular ECs from e12 and e17.5 datasets. (A and B) UMAPs showing expression of selected EC subtype markers in e12 ECs (A) and e17.5 ECs (B). (C and D) Breakdown of cell types as percentage of total cells in e12 (C) and e17.5 (D). (E and F) UMAPs showing the cells that were used for the analysis of e12 coronary ECs in Fig. 1c (E) and for the analysis of e17.5 coronary ECs in Fig. 1f (F). Scale bar from (A) also applies to (B).

**Supplementary Figure 3:**
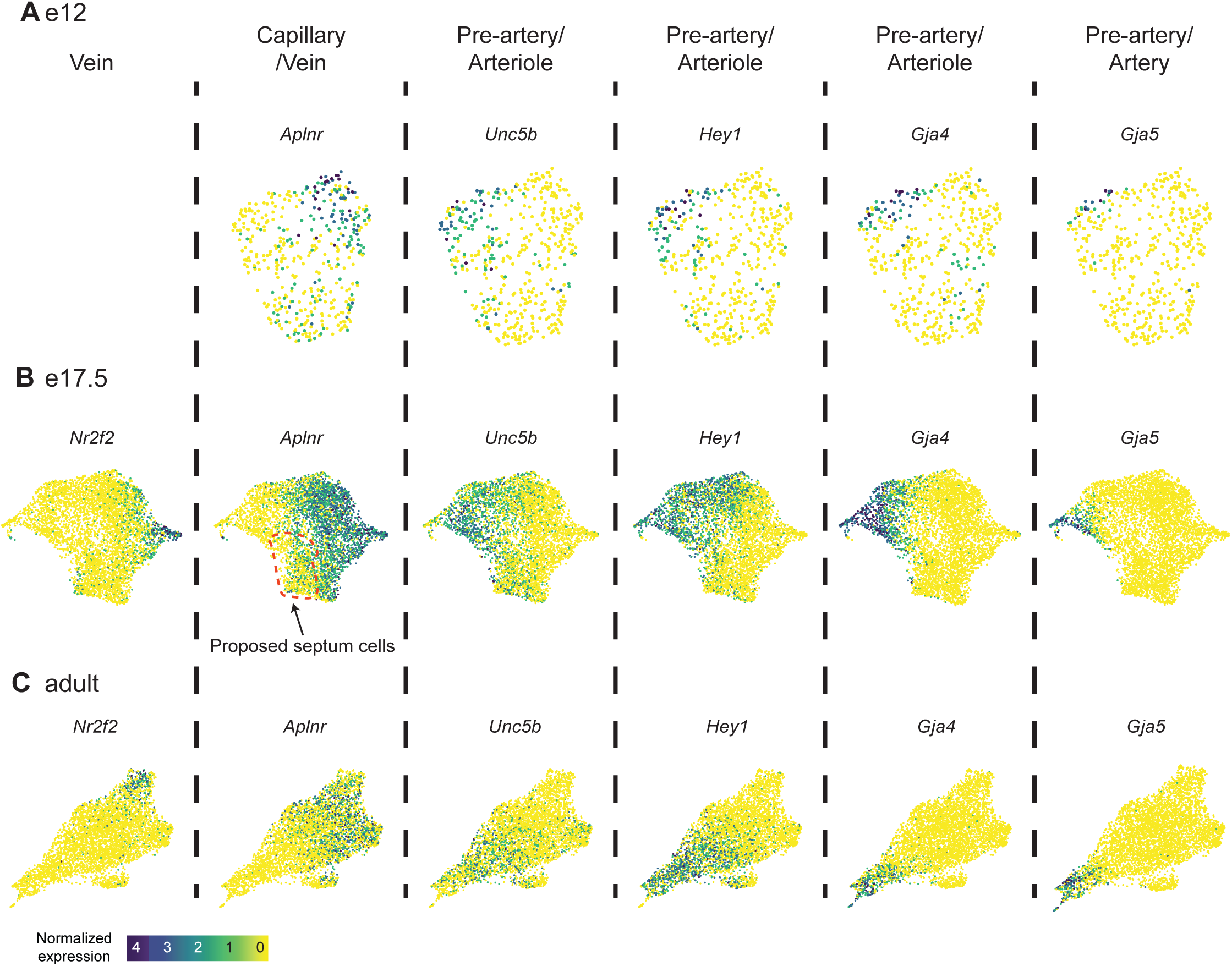
Coronary EC subtype markers. (A, B and C) UMAPs showing expression of selected coronary EC subtype markers in coronary ECs at e12 (A), e17.5 (B) and adult (C). Scale bar from (C) also applies to (A) and (B).

**Supplementary Figure 4:**
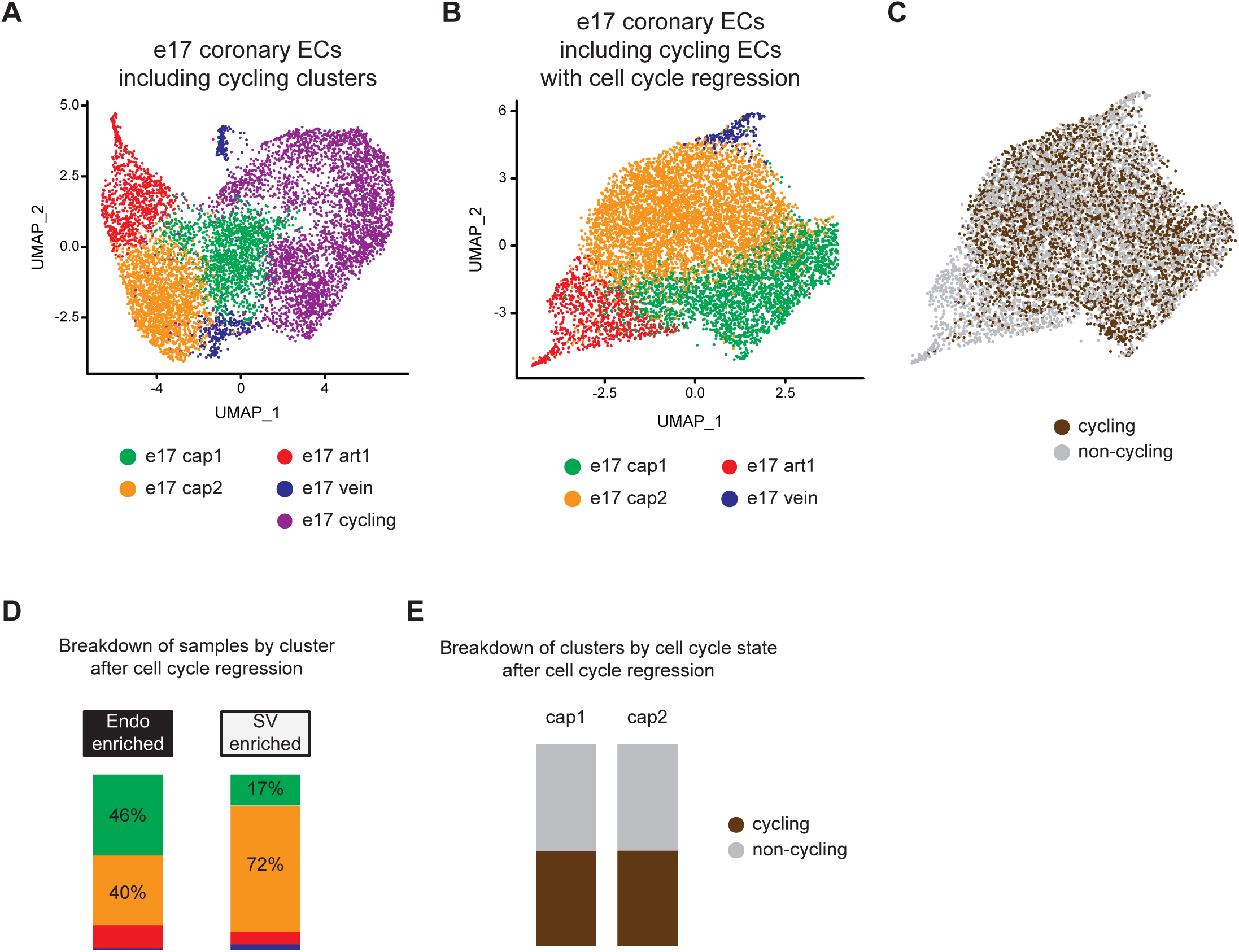
Cell cycle regression in e17.5 coronary ECs. (A) UMAP showing unbiased clustering of e17.5 mouse coronary ECs before the removal of cycling cells. (B) UMAP showing unbiased clustering of e17.5 mouse coronary ECs from (A) after cell cycle regression was performed. (C) Post-regression UMAP from (B) showing the cycling cells which were in the cycling cluster in (A). (D) Breakdown of Endo- and SV-enriched cells from (B) by cluster. (E) Breakdown of the capillary clusters in (B) into cells that are cycling or non-cycling.

**Supplementary Figure 5:**
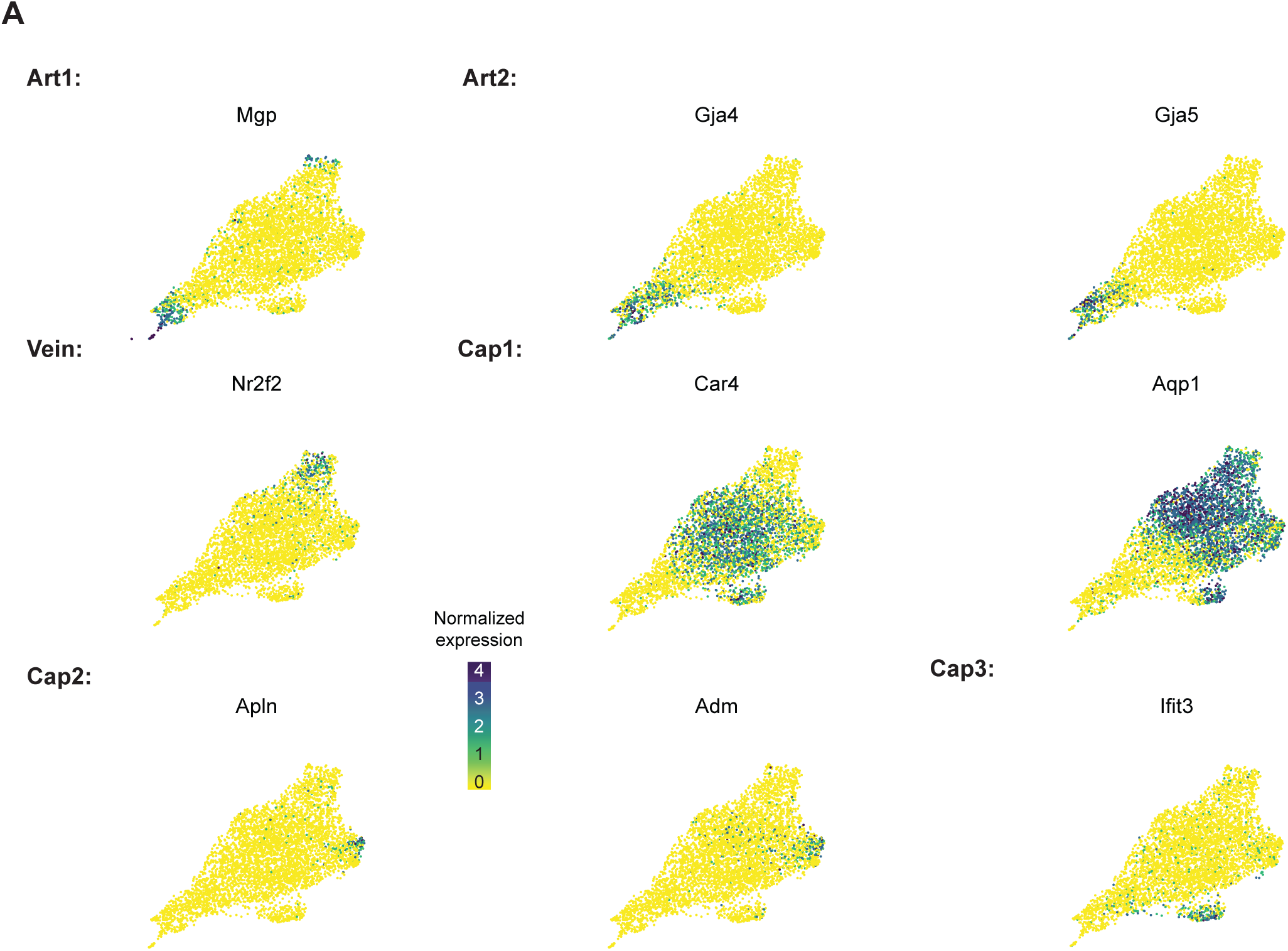
Expression of marker genes adult mouse coronary EC dataset. (A) UMAPs showing expression of selected coronary EC subtype markers in the adult coronary EC dataset from Fig. 1.

**Supplementary Figure 6:**
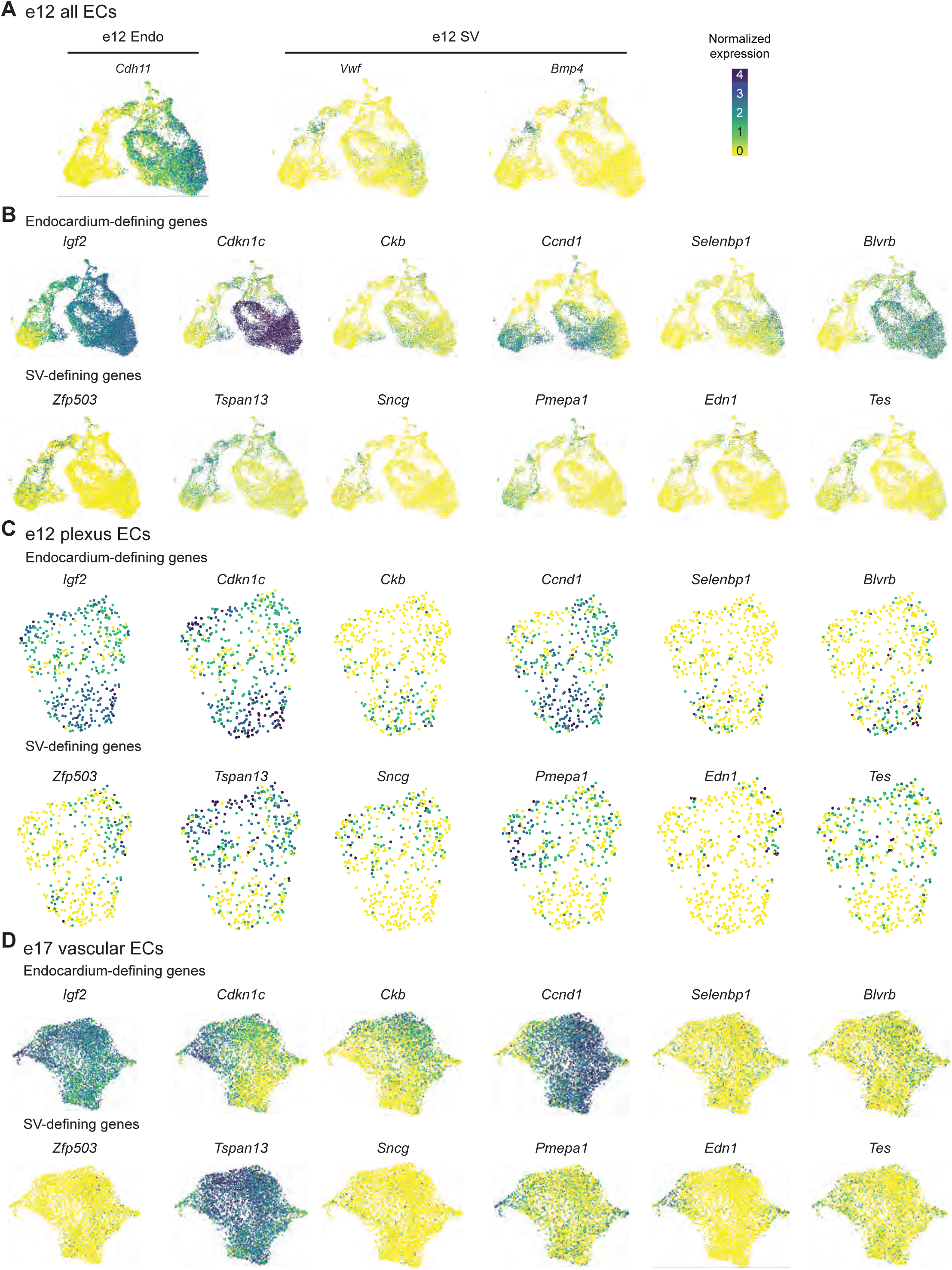
Expression of selected Endo- and SV-defining genes. (A) UMAP showing expression of canonical Endo (*Cdh11*) and SV (*Vwf, Bmp4*) markers in heart ECs at e12. (B, C and D) Expression of genes enriched in either the Endo (Endo-defining genes) or the SV (SV-defining genes) in all e12 ECs (B), e12 coronary plexus ECs (C) and e17.5 coronary ECs (D). Scale bar from (A) also applies to (B), (C), and (D).

**Supplementary Figure 7:**
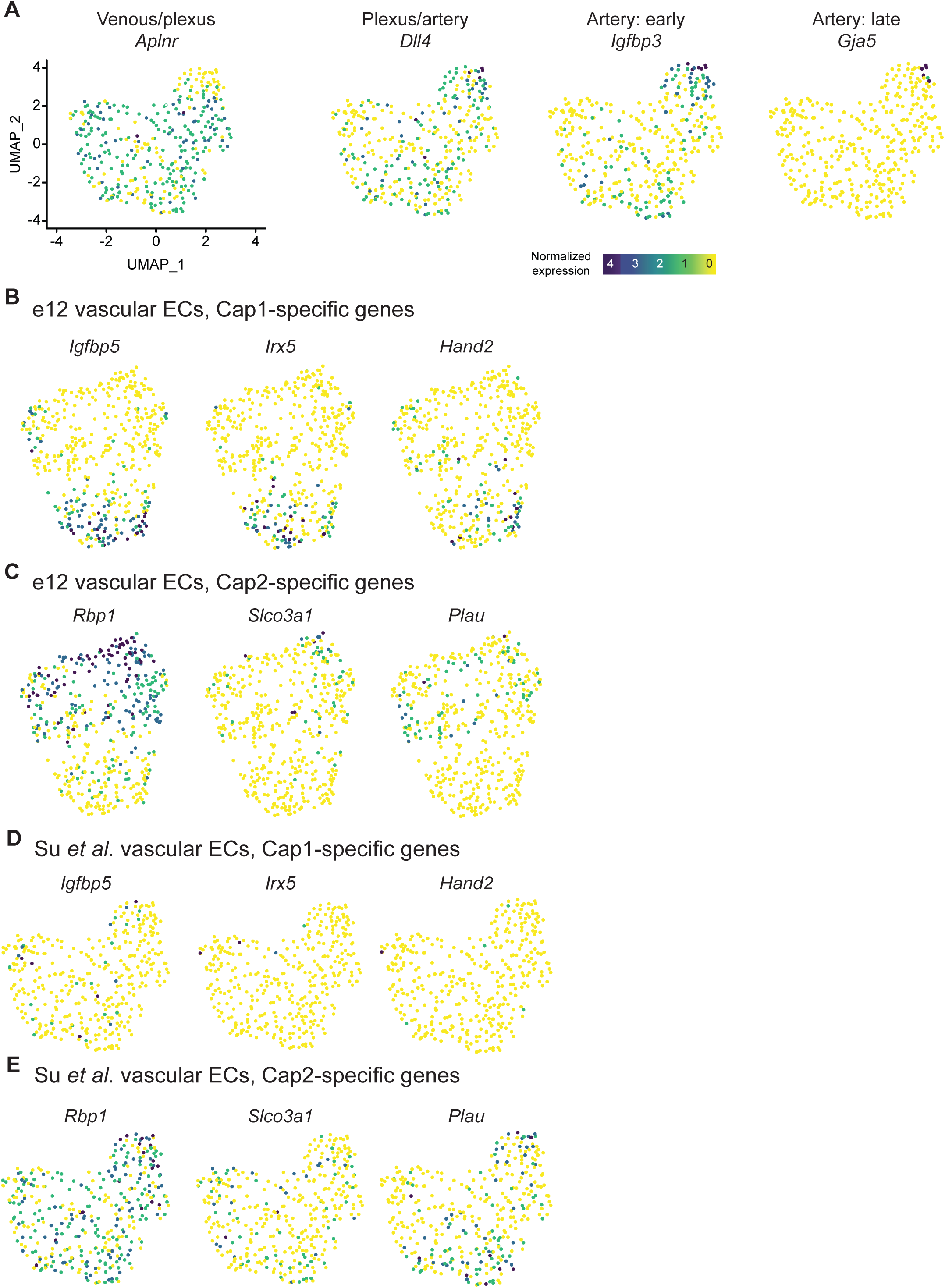
Expression of e12 Cap1- and Cap2-specific genes in a dataset of e12.5 SV-derived ECs. (A) UMAP showing expression of selected coronary EC subtype markers in a previously published dataset (Su et al., 2018). (B and C) Expression in e12 dataset of genes enriched in e12 Cap1 (B) or Cap2 (C). (D and E) Expression in Su *et al*. dataset of genes enriched in e12 Cap1 (D) or Cap2 (E). Scale bar from (A) also applies to (B), (C), (D), and (E).

**Supplementary Figure 8:**
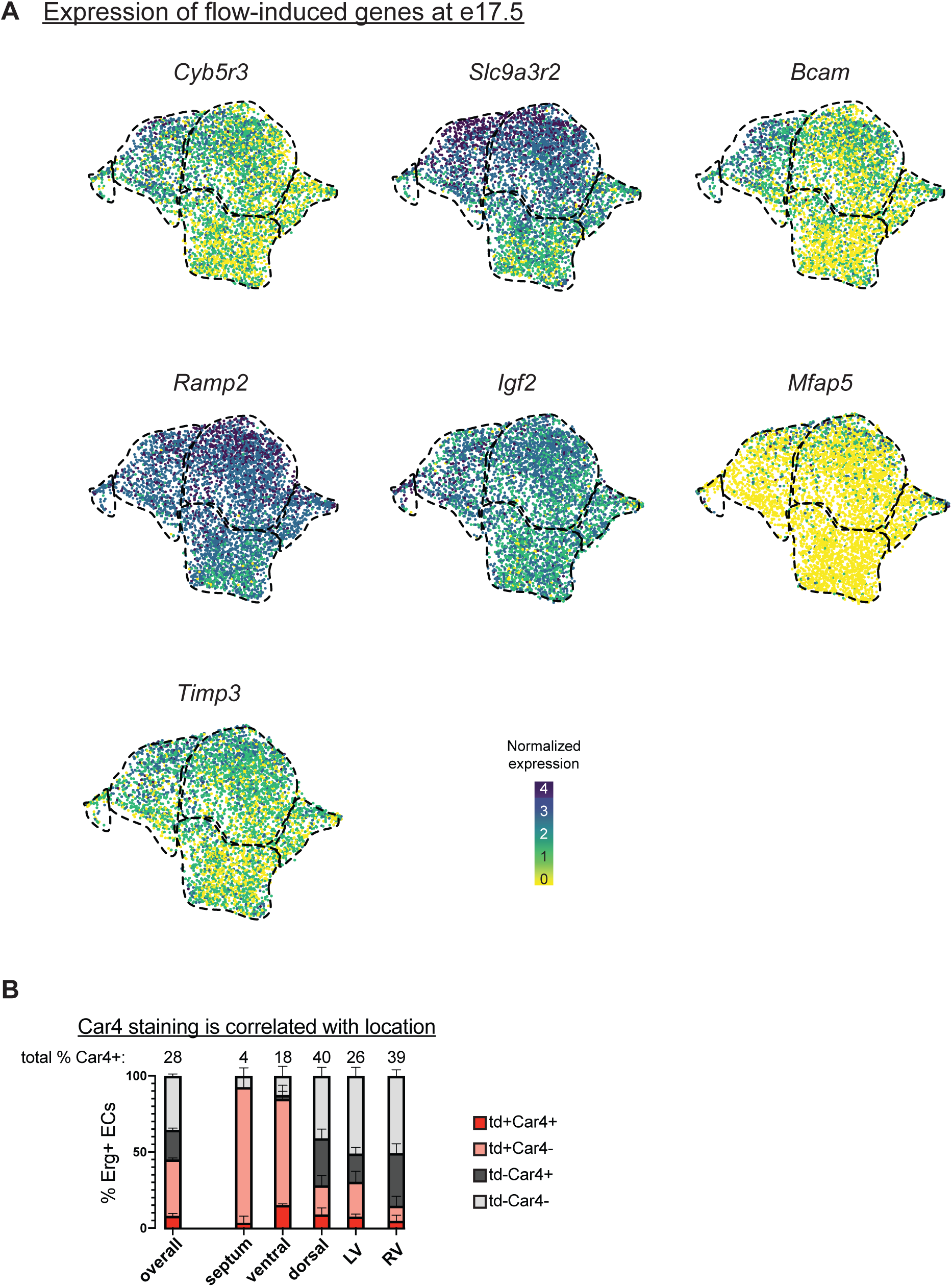
Expression of flow-induced genes. (A) UMAPs showing expression of selected flow-induced genes from Kumar et al 2014 (Kumar et al., 2014) in e17.5 coronary ECs. (B) Quantification of Car4 staining in Erg-positive cells from three e17.5 *BmxCreER;Rosa^tdTomato^* embryos (error bars = SD). Red dashed lines outline the putative septal cells as determined in Fig. 3c.

**Supplementary Figure 9:**
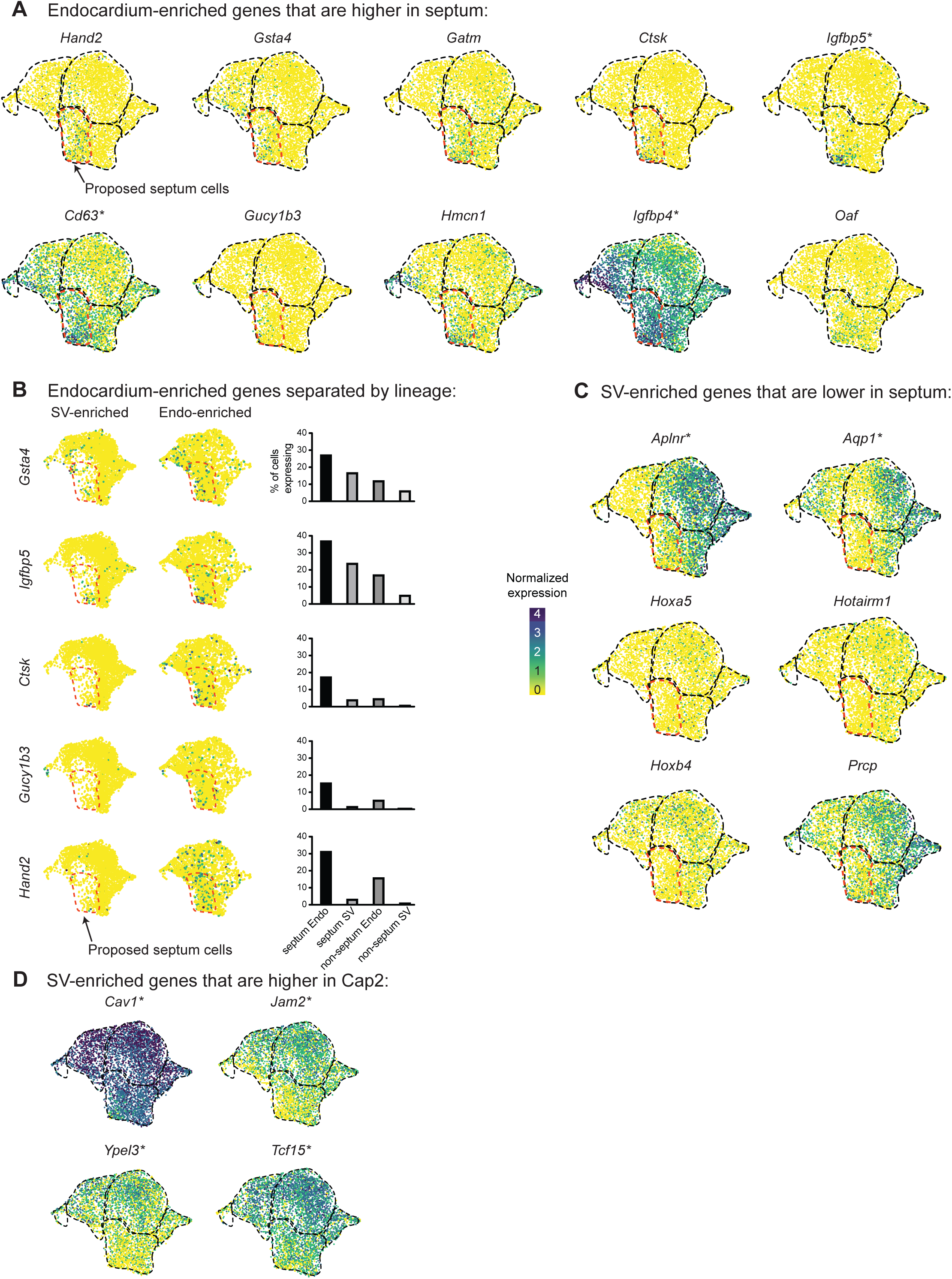
Expression of Endo- and SV-defining genes at e17.5. (A) UMAPs showing expression of Endo-enriched genes manually determined to be expressed in a higher percentage of proposed septum cells (as shown in Fig. 3c) than non-septum cells. (B) UMAPs showing expression of selected Endo-enriched genes split by lineage. Bar plots show the percent of capillary cells in different categories (septum Endo-enriched, septum SV-enriched, non-septum Endo-enriched, non-septum SV-enriched) which express each gene at any level. (C) UMAPs showing expression of SV-enriched genes manually determined to be expressed in a higher percentage of non-septum cells than septum cells. (D) UMAPs showing expression of SV- enriched genes manually determined to be expressed in a higher percentage of Cap2 cells compared to Cap1 cells. Starred genes are significantly differentially expressed between e17.5 Cap1 and Cap2, as indicated in Fig. 2c. Scale bar from (B) also applies to (A), (C), and (D).

**Supplementary Figure 10:**
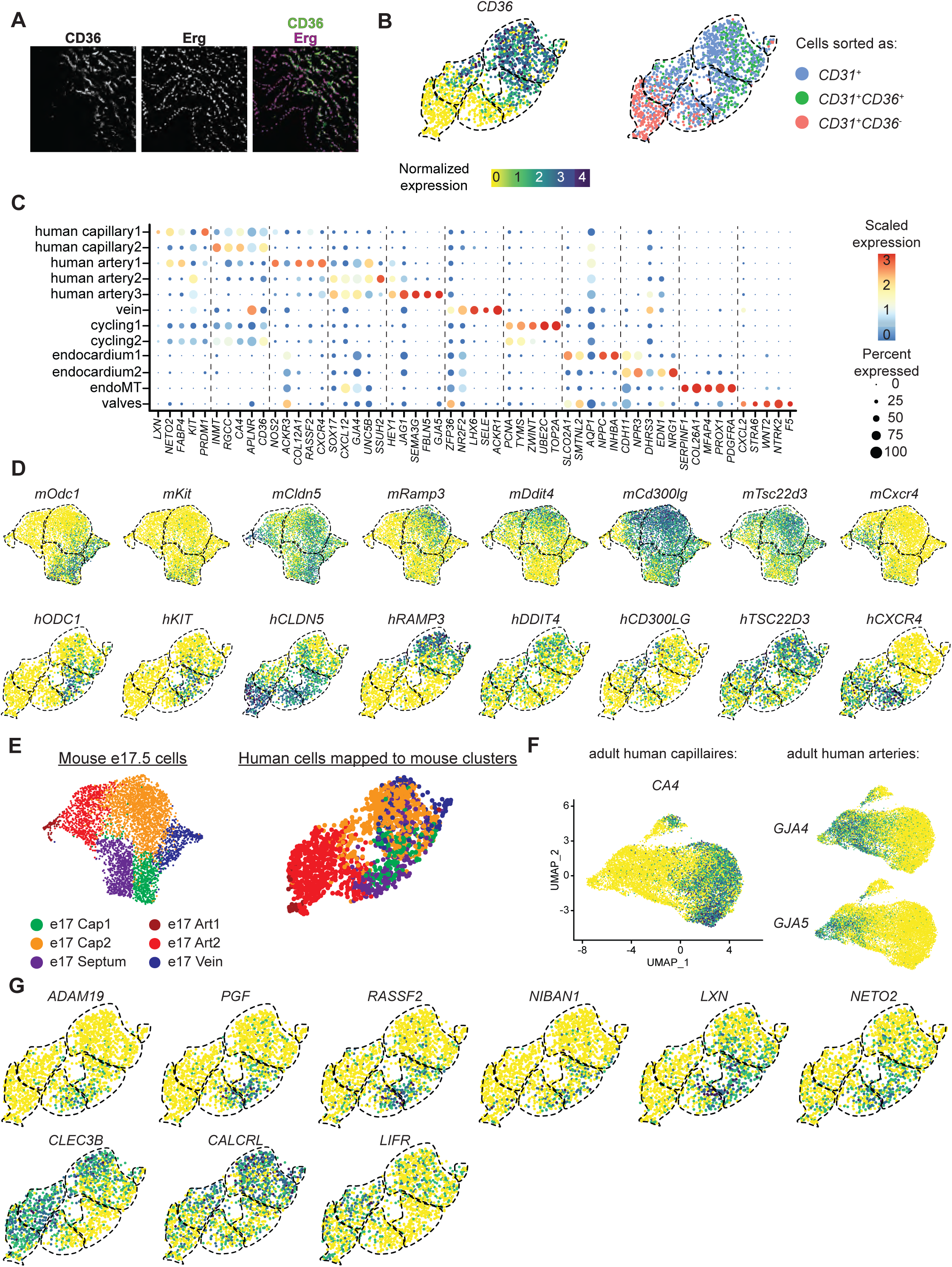
Additional analysis of developing human coronary ECs. (A) Immunofluorescence for *CD36* and *Erg* in a section from a 14 wk human fetal heart. (B) UMAPs showing expression of *CD36* in human coronary ECs as well as cells colored according to FACS sample, i.e. *PECAM1*+ only, *PECAM1+CD36*+, *PECAM1+CD36*-, as indicated in Fig. 5a. Dashed lines show the borders of the previously defined human coronary clusters. (C) Dot plot showing the expression of selected gene markers for each human EC cluster from Fig. 5b. (D) UMAPs showing expression of selected genes with shared expression patterns between mouse e17.5 and human fetal capillary ECs. (E) UMAPs showing mouse e17.5 coronary EC clusters including a manually-defined septum cluster as shown in Fig. 3c, and the fetal human coronary ECs which map to each of these clusters. (F) UMAPs showing expression of selected capillary and artery genes in adult human coronary ECs from a previously published dataset (Litvinukova et al., 2020). (G) UMAPs showing expression of selected genes shared between human Cap1 and Art3 or between human Cap2 and Art2. Scale bar from (B) also applies to (D), (F), and (G).

**Supplementary Figure 11:**
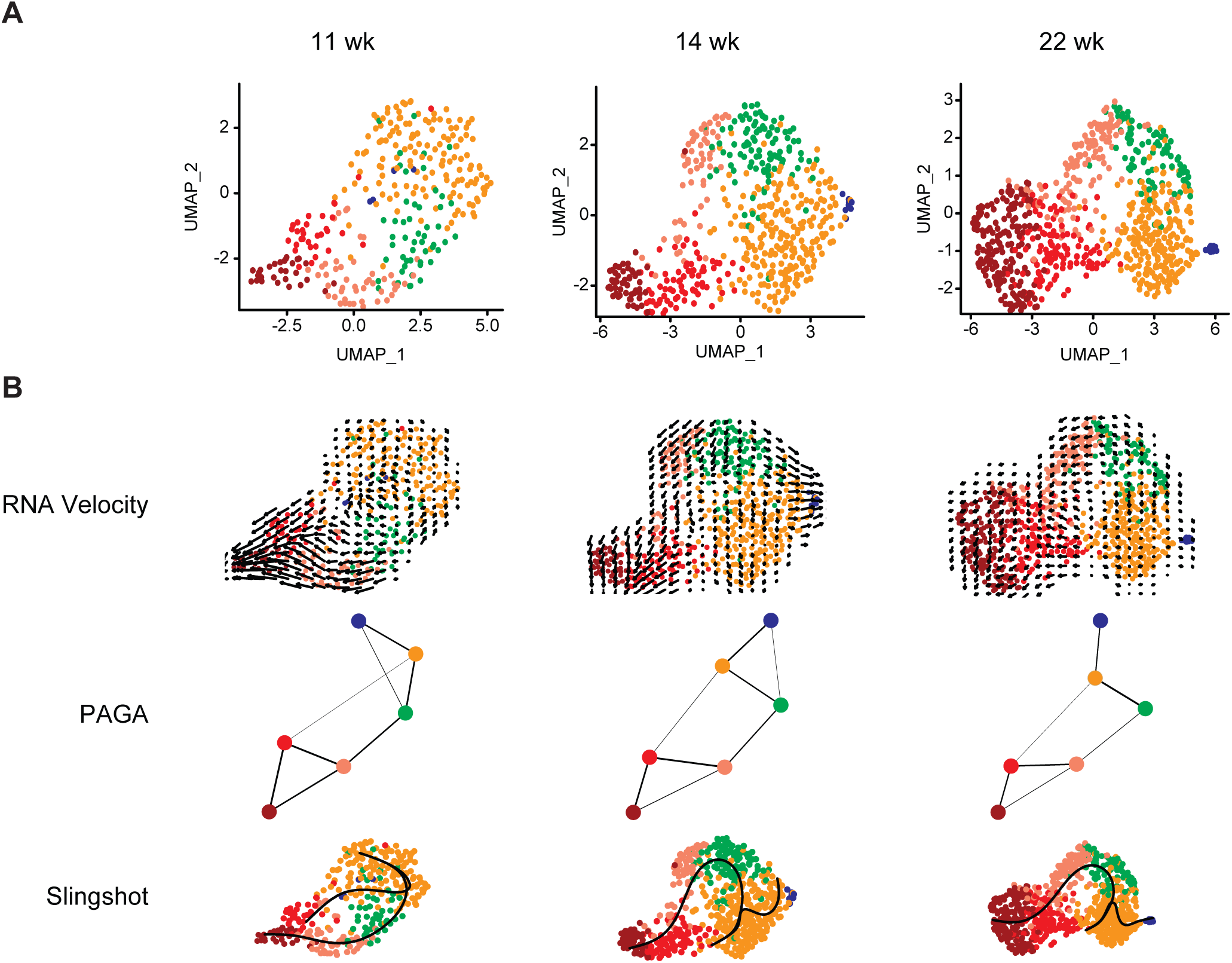
Analysis of developing human coronary ECs separated by stage. (A) UMAPs showing unbiased clustering of cells isolated from each individual human fetal heart. (B) Trajectory analysis of human coronary EC at each individual stage using RNA velocity, PAGA, Slingshot, and Monocle.

